# Host-pathogen genetic interactions underlie tuberculosis susceptibility

**DOI:** 10.1101/2020.12.01.405514

**Authors:** Clare M. Smith, Richard E. Baker, Megan K. Proulx, Bibhuti B. Mishra, Jarukit E. Long, Sae Woong Park, Ha-Na Lee, Michael C. Kiritsy, Michelle M. Bellerose, Andrew J. Olive, Kenan C. Murphy, Kadamba Papavinasasundaram, Frederick J. Boehm, Charlotte J. Reames, Rachel K. Meade, Brea K. Hampton, Colton L. Linnertz, Ginger D. Shaw, Pablo Hock, Timothy A. Bell, Sabine Ehrt, Dirk Schnappinger, Fernando Pardo-Manuel de Villena, Martin T. Ferris, Thomas R. Ioerger, Christopher M. Sassetti

## Abstract

The outcome of an encounter with *Mycobacterium tuberculosis* depends on the pathogen’s ability to adapt to the variable immune pressures exerted by the host. Understanding this interplay has proven difficult, largely because experimentally tractable animal models do not recapitulate the heterogeneity of tuberculosis disease. We leveraged the genetically diverse Collaborative Cross (CC) mouse panel in conjunction with a library of *Mtb* mutants to associate bacterial genetic requirements with host genetics and immunity. We report that CC strains vary dramatically in their susceptibility to infection and produce qualitatively distinct immune states. Global analysis of *Mtb* mutant fitness across the CC panel revealed that many virulence pathways are only in specific host microenvironments, identifying the large fraction of the pathogen’s genome that has been maintained to ensure fitness in a diverse population. Both immunological and bacterial traits were associated with genetic variants distributed across the mouse genome, identifying the specific host-pathogen genetic interactions that influence pathogenesis.

## Introduction

Infection with *Mycobacterium tuberculosis* (*Mtb*) produces heterogeneous outcomes that are influenced by genetic and phenotypic variation in both the host and the pathogen. Classic human genetic studies show that host variation influences immunity to tuberculosis (TB) (Abel et al., 2018; Comstock, 1978). Likewise, the co-evolution of *Mtb* with different populations across the globe has produced genetically distinct lineages that demonstrate variable virulence traits (Gagneux et al., 2006). The role of genetic variation on each side of this interaction is established, yet the intimate evolutionary history of both genomes suggests that interactions between host and pathogen variants may represent an additional determinant of outcome (McHenry et al., 2020). Evidence for genetic interactions between host and pathogen genomes have been identified in several infections (Ansari et al., 2017; Berthenet et al., 2018), including TB (Caws et al., 2008; Holt et al., 2018; Thuong et al., 2016). However, the combinatorial complexity involved in identifying these relationships in natural populations have left the mechanisms largely unclear.

Mouse models have proven to be a powerful tool to understand mechanisms of susceptibility to TB. Host requirements for protective immunity were discovered by engineering mutations in the genome of standard laboratory strains of mice, such as C57BL/6 (B6), revealing a critical role of Th1 immunity. Mice lacking factors necessary for the production of Th1 cells or the protective cytokine interferon gamma (IFNγ) are profoundly susceptible to *Mtb* infection (Caruso et al., 1999; Cooper et al., 1993; Cooper et al., 1997; Flynn et al., 1993; Saunders et al., 2002). Defects in this same immune axis cause the human syndrome Mendelian Susceptibility to Mycobacterial Disease (MSMD) (Altare et al., 1998; Bogunovic et al., 2012; Bustamante et al., 2014; Filipe-Santos et al., 2006), demonstrating the value of knockout (KO) mice to characterize genetic variants of large effect. Similarly, the standard mouse model has been used to define *Mtb* genes that are specifically required for optimal bacterial fitness during infection (Bellerose et al., 2020; Sassetti and Rubin, 2003; Zhang et al., 2013).

Despite the utility of standard mouse models, it has become increasingly clear that the immune response to *Mtb* in genetically diverse populations is more heterogeneous than any single small animal model (Smith and Sassetti, 2018). For example, while IFNγ-producing T cells are critical for protective immunity in standard inbred lines of mice, a significant fraction of humans exposed to *Mtb* control the infection without producing a durable IFNγ response (Lu et al., 2019). Similarly, IL-17 producing T cells have been implicated in both protective responses and inflammatory tissue damage in TB, but IL-17 has little effect on disease progression in B6 mice, except in the context of vaccination or infection with particularly virulent *Mtb* (Gopal et al., 2012; Khader et al., 2007). The immunological homogeneity of standard mouse models may also explain why only a small minority of the >4000 genes that have been retained in the genome of *Mtb* during its natural history promote fitness in the mouse (Bellerose et al., 2020). Thus, homogenous mouse models of TB fail to capture the distinct disease states, mechanisms of protective immunity, and selective pressures on the bacterium that are observed in natural populations.

The Collaborative Cross (CC) and Diversity Outbred (DO) mouse populations are new mammalian resources that more accurately represent the genetic and phenotypic heterogeneity observed in outbred populations (Churchill et al., 2004; Churchill et al., 2012). These mouse panels are both derived from the same eight diverse founder strains but have distinct population structures (Saul et al., 2019). DO mice are maintained as an outbred population and each animal represents a unique and largely heterozygous genome (Keller et al., 2018; Svenson et al., 2012). In contrast, each inbred CC strain’s genome is almost entirely homozygous, producing a genetically stable and reproducible population in which the phenotypic effect of recessive mutations is maximized (Shorter et al., 2019; Srivastava et al., 2017). Together, these resources have been leveraged to identify host loci underlying the immune response to infectious diseases (Noll et al., 2019). In the context of TB, DO mice have been used as individual, unique hosts to identify correlates of disease, which resemble those observed in non-human primates and humans (Ahmed et al., 2020; Gopal et al., 2013; Niazi et al., 2015). Small panels of the reproducible CC strains have been leveraged to identify host background as a determinant of the protective efficacy of BCG vaccination (Smith et al., 2016) and a specific variant underlying protective immunity to tuberculosis (Smith et al., 2019). While these studies demonstrate the tractability of the DO and CC populations to model the influence of host diversity on infection, dissecting host-pathogen interactions requires the integration of pathogen genetic diversity.

We combined the natural but reproducible host variation of the CC panel with a comprehensive library of *Mtb* mutants to characterize the interactions between host and pathogen. Using over 60 diverse mouse strains, we report that the CC panel encompasses a broad spectrum of TB susceptibility and immune phenotypes, including outlier lines that model non-canonical immune states. Through “Transposon Sequencing” (TnSeq), we quantified the relative fitness of *Mtb* mutants across the CC panel and specific immunological knockout strains. We report that approximately three times more bacterial genes contribute to fitness in the diverse panel than in any single mouse strain, defining a large fraction of the bacterial genome that is dedicated to adapting to distinct immune states. Association of these immunological and bacterial fitness traits with Quantitative Trait Loci (QTL) demonstrated the presence of discrete Host-Interacting-with Pathogen QTL (*Hip*QTL) that represent inter-species genetic interactions that influence the pathogenesis of this infection.

## Results

### The spectrum of TB disease traits in the CC exceeds that observed in standard inbred mice

To characterize the diversity of disease states that are possible in a genetically diverse mouse population, we infected a panel of 52 CC lines and the 8 founder strains with *Mtb*. To enable TnSeq studies, the animals were infected via the intravenous route with a saturated library of *Mtb* transposon mutants, which in sum produce an infection that is similar to the wild-type parental strain (Bellerose et al., 2020; Sassetti and Rubin, 2003). Groups of 2-6 (average of n=3) male mice per genotype were infected, and the bacterial burden after four weeks of infection was assessed by plating (colony forming units, CFU) and quantifying the number of bacterial chromosomes in the tissue (chromosome equivalents, CEQ). These two metrics were highly correlated (r=0.88) and revealed a wide variation in bacterial burden across the panel (**Figure 1A** and **S1**). The susceptibility of the inbred founder strains was largely consistent with previous studies employing an aerosol infection (Smith et al., 2016). As expected, B6 mice were relatively resistant to infection, along with 129S1/SvlmJ (129) and NOD/ShiLtJ (NOD) strains. In contrast to the standard inbred lines, lung bacterial burden varied by more than 1000-fold across the more diverse CC panel, ranging from animals that are significantly more resistant than B6, to mice that harbored more than 10^9^ bacteria in their lungs (**Figure 1A**). Bacterial burden in the spleen also varied several thousand-fold across the panel and was moderately correlated with lung burden (r=0.43) (**Table S1** and **Figure S1**). Thus, the CC panel encompasses a much greater quantitative range of susceptibility than standard inbred lines.

**Figure 1.**
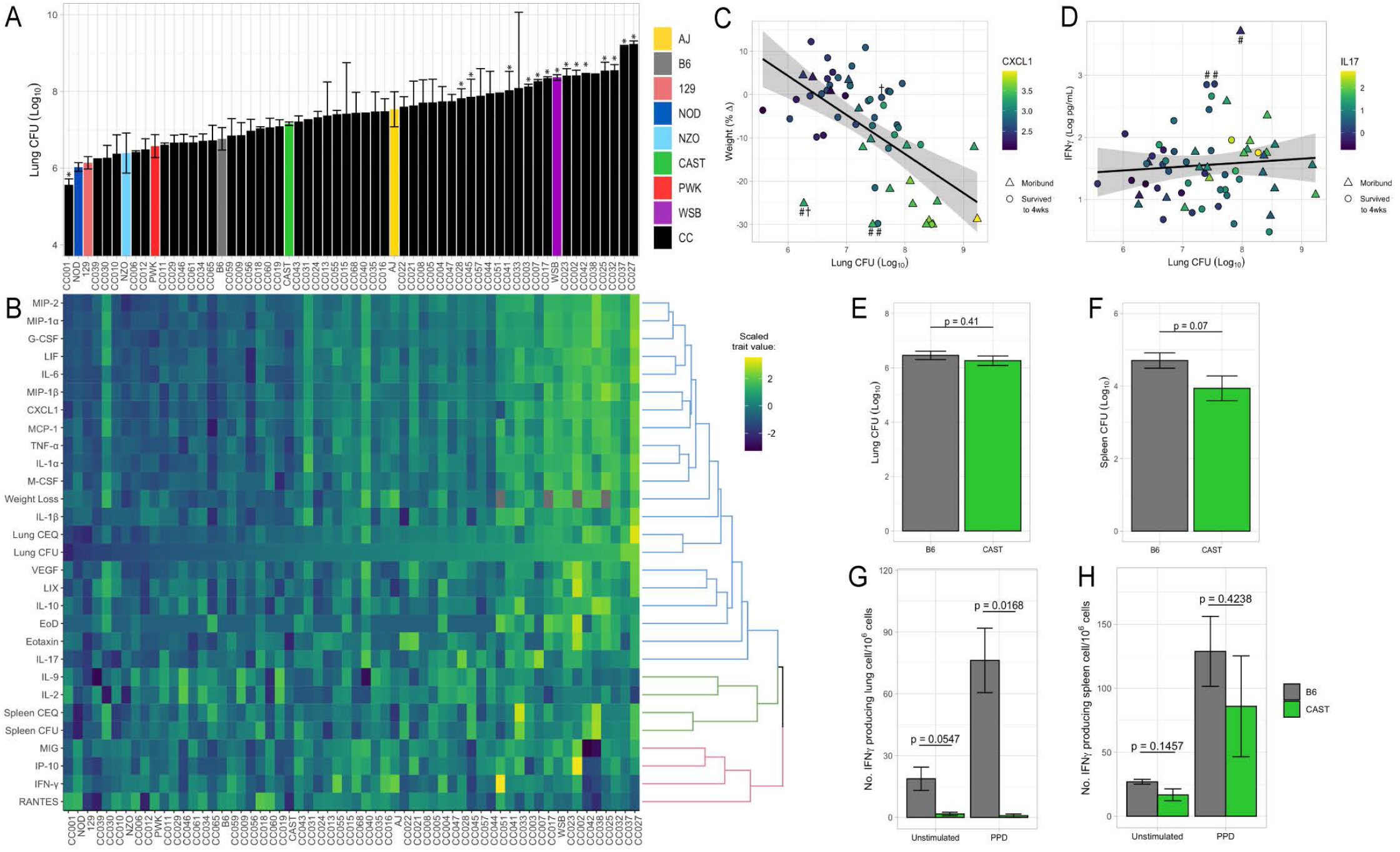
The spectrum of M. tuberculosis disease-related traits across the Collaborative Cross. (A) Average lung CFU (log10) across the CC panel at 4 weeks post-infection. Bars show mean+/-SD for CFU per CC or parental strain; groups of 2-6 (average of n=3) mice per genotype. Bars noted with ∗ indicate strains that were statistically different from B6 (P<0.05; 1-factor ANOVA with Dunnett’s post-test). (B) Heatmap of the 32 disease-related traits (log10 transformed) measured including: lung and spleen colony forming units (CFU); lung and spleen chromosomal equivalents (CEQ); weight loss(% change); cytokines from lung, “earliness of death” (EoD) reflecting the number of days prior to the end of experiment that moribund strains were euthanized. Mouse genotypes are ordered by lung CFU. Scaled trait values were clustered (hclust in R package heatmaply) and dendrogram nodes colored by 3 k-means. Blue node reflects correlation coefficient r>0.7; green r=0.3-0.6 and red r<0.2. (C) Correlation of lung CFU and weight(% change) shaded by CXCL1 levels. Genotypes identified as statistical outliers for weight are noted by#; CXCL1 by †. (D) Correlation of lung CFU and IFNγ levels shaded by IL-17. Strains identified as outliers for IFNγ noted by #. Each point in (C) and (D) is the average value per genotype. Outlier genotypes were identified after linear regression using studentized residuals. (E-H) B6 and CAST traits measured at 4 weeks post aerosol infection (E) lung CFU; (F) spleen CFU; (G) number of IFNγ producing lung T cells; (H) number of IFNγ producing spleen T cells. Bar plots show the mean +/-SD. Significance between groups was determined by Welch’s t test. Groups consist of 4-10 mice per genotype from 2 independent experiments. All mice in the initial CC screen were male; validation studies in panels E-H included both sexes, where no statistical difference at the 4-week timepoint between sexes was observed.

Comparing various measures of infection progression showed many expected correlations but also an unexpected decoupling of some phenotypes. As an initial assessment of the disease processes in these animals, we correlated bacterial burden and lung cytokine abundance with measures of systemic disease such as weight loss and sufficient morbidity to require euthanasia (“earliness of death”). In general, correlations between these metrics indicated that systemic disease was associated with bacterial replication and inflammation (**Figure 1B** and **Figure S1**). Lung CFU was correlated with weight loss, mediators that enhance neutrophil differentiation or migration (CXCL2 (MIP-2; r=0.79), CCL3 (MIP-1a; r=0.77), G-CSF (r=0.78) and CXCL1 (KC; r=0.76)), and more general proinflammatory cytokines (IL-6 (r=0.80) and IL-1α (r=0.76)) (**Figure S1)**. These findings are consistent with previous work in the DO panel, that found both proinflammatory chemokines and neutrophil accumulation to be predictors of disease (Ahmed et al., 2020; Gopal et al., 2013; Niazi et al., 2015).

The reproducibility of CC genotypes allowed us to quantitatively assess the heritability (h^2^) of these immunological and disease traits. The percent of the variation attributed to genotype ranged from 56-87% (mean=73.4%; **Table S2**). The dominant role of genetic background in determining the observed phenotypic range allowed a rigorous assessment of outlier phenotypes that is not possible in the DO population. For example, despite the correlation between lung CFU and weight loss (r=0.57), several strains failed to conform to this relationship (**Figure 1C**). In particular, CC030/GeniUnc (p=0.003), CC040/TauUnc (p=0.027) and A/J (p=0.03) lost significantly more weight than their bacterial burdens would predict (**Figure 1C**; outlier genotypes determined by studentized residuals; noted by #). Similarly, CXCL1 abundance was significantly higher in CC030/GeniUnc (p=0.001) and lower in CC056/GeniUnc (p=0.040), than the level predicted by their respective bacterial burden (**Figure 1C**; outlier genotypes noted by †). Thus, these related disease traits can be dissociated based on the genetic composition of the host.

The cluster of cytokines that was most notably unrelated to bacterial burden included IFNγ and the interferon-inducible chemokines CXCL10 (IP10), CXCL9 (MIG), and CCL5 (RANTES) (Red cluster in **Figure 1B**; **Figure S1**) (r<0.3). Despite the clear protective role for IFNγ (Cooper et al., 1993; Flynn et al., 1993), high levels have been observed in susceptible mice, likely as a result of high antigen load (Barber et al., 2011; Lázár-Molnár et al., 2010). While high IFNγ levels in susceptible animals was therefore expected, it was more surprising to find a number of genotypes that were able to control bacterial replication yet had very low levels of this critically important cytokine (**Figure 1D**). This observation is likely due the inclusion of two founder lines, CAST/EiJ (CAST) and PWK/PhJ (PWK) that display this unusual phenotype (Smith et al., 2016). To further investigate, a separate cohort of B6 and CAST animals was infected by the aerosol route, and the number of IFNγ-producing T cells in lung and spleen was compared by ELISPOT assays. At 4 weeks post-infection, B6 and CAST harbored comparable burdens of *Mtb* in lung and spleen (**Figure 1E** and **1F**), and the infection elicited similar numbers of *Mtb*-specific IFNγ-producing cells in the spleen (**Figure 1H**). In contrast, while IFNγ producing cells were found in the lungs of B6 mice, none were detectable in CAST (**Figure 1G**). Thus, while CAST animals are capable of producing IFNγ-secreting cells in response to *Mtb* infection, these cells do not appear to be involved in bacterial control in the lung. As CC strains that share this immune profile also produced low levels of IL-17 (**Figure 1D**), the mechanism(s) conferring protection in these animals remain unclear. In sum, this survey of TB-related traits in the CC demonstrated a broad range of susceptibility and the presence of qualitatively distinct and genetically determined disease states.

### *Tip*QTL define genetic variants that control TB immunophenotypes

Tuberculosis ImmunoPhenotype Quantitative Trait Loci (*Tip*QTL), which were associated with TB disease or cytokine traits, were identified and numbered in accordance with previously reported *Tip*QTL (Smith et al., 2019). Of the 32 TB-disease traits, we identified 9 individual metrics that were associated with reasonable statistical confidence to a chromosomal locus. Of these, three were associated with high confidence (p≤0.053), and six other QTL met a suggestive threshold (p<0.2; **Table 1**). Several individual trait QTL occupied the same chromosomal locations. For example, spleen CFU and spleen CEQ, which are both measures of bacterial burden and highly correlated, were associated with the same interval on distal chromosome 2 (**Table 1**, *Tip5*; **Figure 2A** and **2C**). IL-10 abundance was associated with two distinct QTL (**Table 1**). While IL-10 was relatively uncorrelated with spleen CFU (r=0.48), one of its QTL fell within the *Tip5* bacterial burden interval on chromosome 2 (**Figure 2A** and **2C**). At this QTL, the NOD haplotype was associated with high values for all three traits (**Figure 2E**). Similarly, the correlated traits, CXCL1 abundance and lung CFU, were individually associated to the same region on chromosome 7 (**Table 1**, *Tip8*; **Figure 2B** and **2D**). In this interval, the CAST haplotype was associated with both low bacterial burden and CXCL1 (**Figure 2F**). At both *Tip5* and *Tip8*, we found no statistical evidence that the positions of the associated QTL were different (*Tip5* p=0.55; *Tip8* p=0.27; 400 bootstrap samples)(Boehm et al., 2019). These observations support the role of a single causal variant at each locus that is responsible for a pleiotropic trait. Coincident mapping can provide both additional statistical support for QTL (p values by Fisher’s combined probability test: Chr 7, p=0.067; Chr 2, p=0.041) and suggests potential mechanisms of disease progression.

**Table 1.** Disease-related Tuberculosis ImmunoPhenotype QTL (*Tip*QTL). Multiple QTL within the same interval and clear allele effects are designated with the same *Tip*QTL number. Column headings: QTL, quantitative trait loci; Chr, chromosome; LOD, logarithm of the odds; CEQ, chromosomal equivalents.

| QTL | Trait | Chr | LOD | P value | Interval start (Mb) | Peak (Mb) | Interval end (Mb) |
| --- | --- | --- | --- | --- | --- | --- | --- |
| ***Tip5*** | Spleen CEQ | 2 | 9.14 | 2.38E-02 | 174.29 | 178.25 | 178.25 |
| ***Tip5*** | Spleen CFU | 2 | 7.04 | 2.19E-01 | 73.98 | 174.29 | 180.10 |
| ***Tip6*** | IL-9 | 2 | 8.61 | 4.52E-02 | 33.43 | 41.4 | 41.48 |
| ***Tip6*** | IL-9 | 2 | 7.85 | 1.26E-01 | 22.77 | 24.62 | 25.65 |
| ***Tip7*** | IL-17 | 15 | 7.84 | 5.27E-02 | 67.98 | 74.14 | 82.11 |
| ***Tip8*** | CXCL1 | 7 | 7.57 | 1.06E-01 | 30.43 | 45.22 | 46.72 |
| ***Tip8*** | Lung CFU | 7 | 7.47 | 1.17E-01 | 31.06 | 37.78 | 45.22 |
| ***Tip9*** | IL-10 | 17 | 7.16 | 1.85E-01 | 80.98 | 82.47 | 83.55 |
| ***Tip10*** | Lung CFU | 15 | 7.13 | 1.86E-01 | 77.00 | 78.16 | 78.70 |

**Figure 2.**
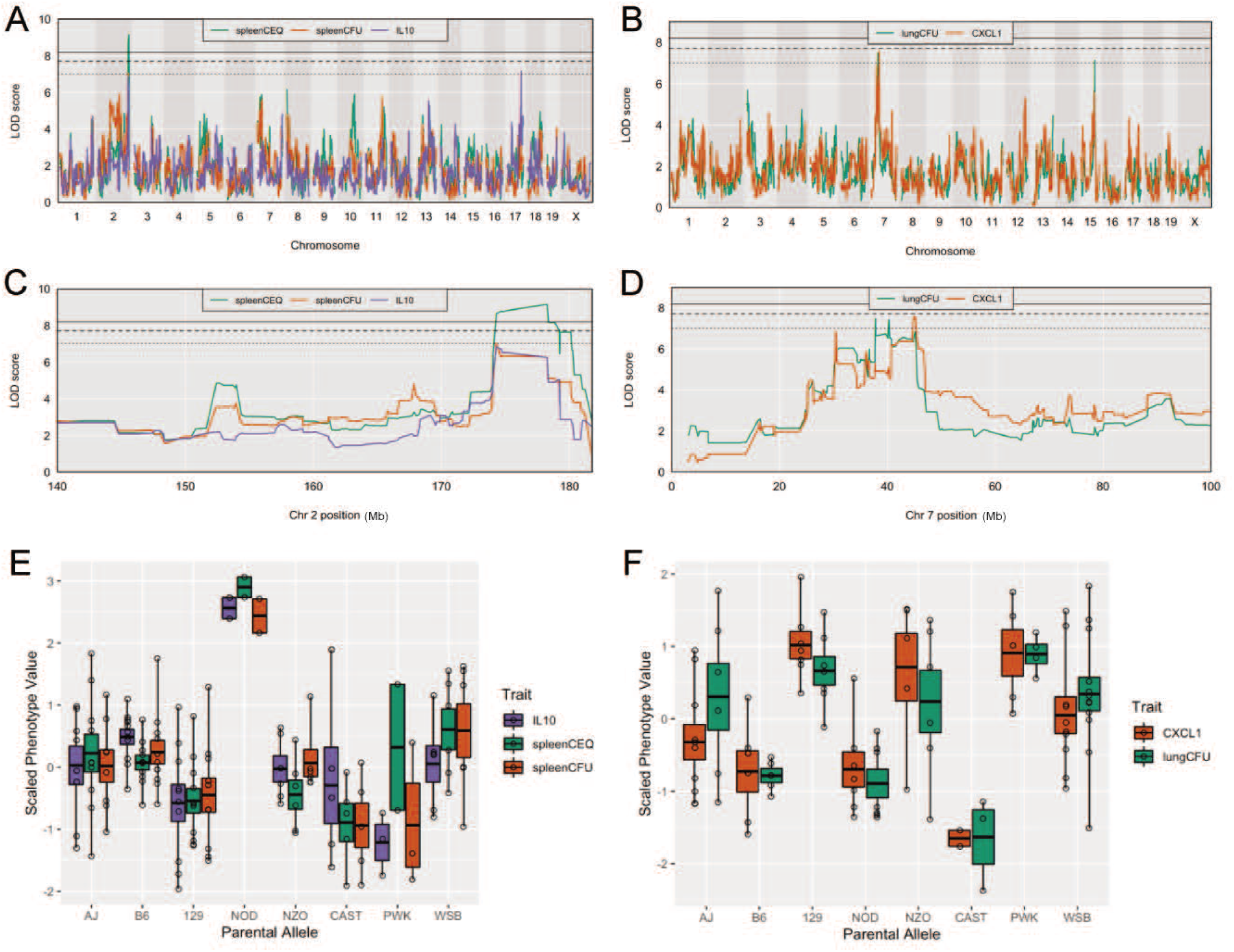
Host loci underlying TB disease-related traits. (A-B) Whole genome QTL scans of (A) spleen CEQ, spleen CFU and IL-10 (B) lung CFU and CXCL1. (C) Zoom of chromosome 2 loci. (D) Zoom of chromosome 7 loci. Thresholds were determined by permutation analysis; solid line, middle dashed line, and lowest dotted lines represent P = 0.05, P = 0.1, and P = 0.2. (E-F) Scaled phenotype value per haplotype at the QTL peak marker. Each dot represents the mean value for a genotype.

A number of factors can limit the statistical significance of QTL identified in the CC population, including small effect sizes, limited genotype availability, and the genetic complexity of the trait. To determine if the statistically suggestive QTL were likely to reflect genuine biological effects, we took an intercross approach to validate the lung CFU QTL on chromosomes 7 and 15 (*Tip8* and *Tip10*, **Table 1**). Given that the associations at both QTL were driven by the CAST haplotype (**Figure 2F**), we generated an F_2_ population based on two CC strains, CC029/Unc and CC030/GeniUnc, that contained CAST sequence at *Tip8* and *Tip10*, respectively (**Figure 3A**). 46 F_2_ mice that were homozygous at either locus were infected with *Mtb*, and lung CFU were enumerated as per the larger CC screen (**Table S3**). Compared to F_2_ mice that were not CAST at either locus, mice that contained CAST at *Tip8, Tip10* or both loci had reduced CFU burden; however, the strongest predictor of significantly lower bacterial burden was CAST at *Tip8* (p=0.005, 2-factor regression; **Figure 3B**). This study provides strong independent support for a CAST-driven QTL on chromosome 7 that controls lung CFU and verifies the validity of the genetic mapping in this dataset.

**Figure 3.**
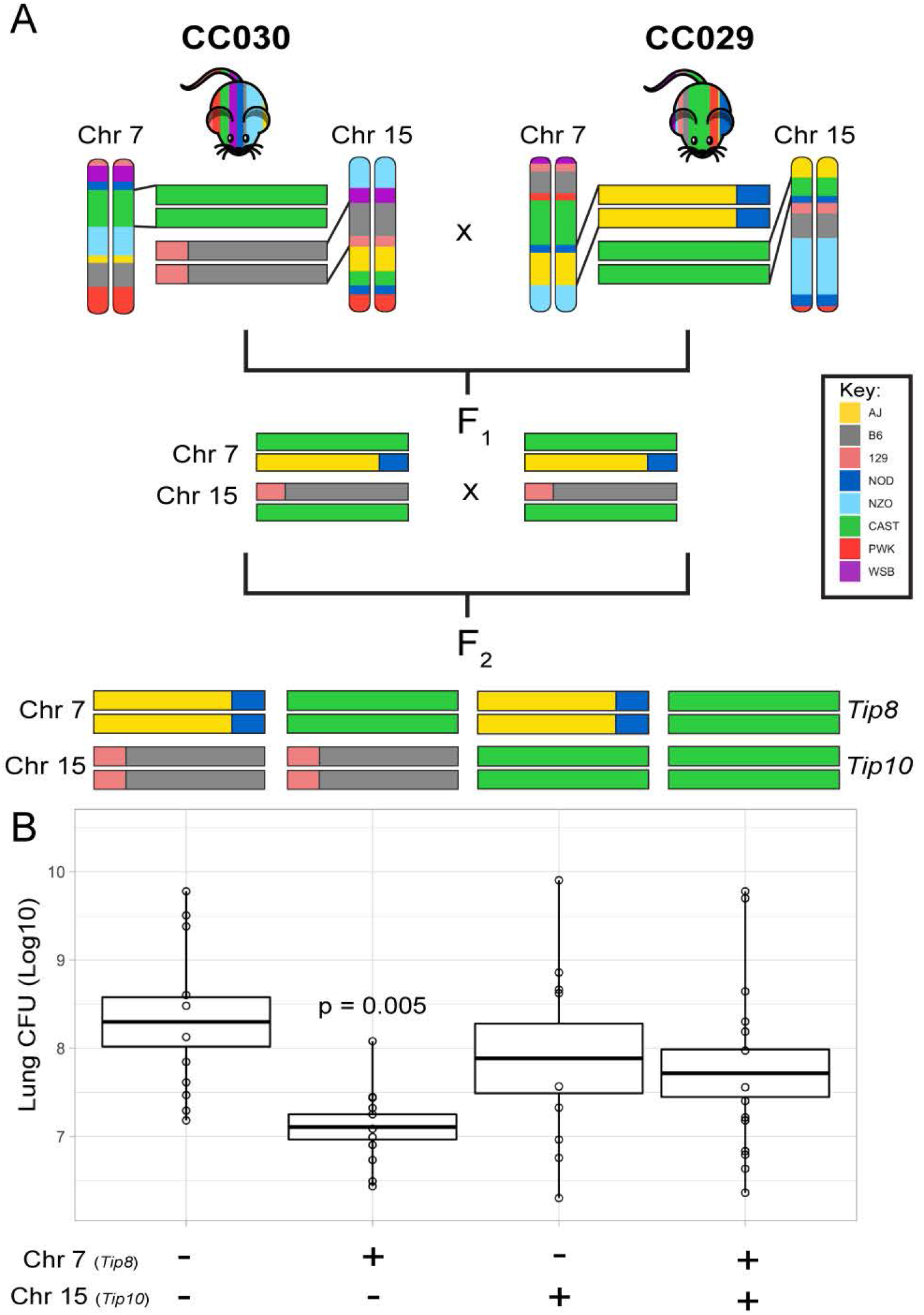
An F2 intercross approach to validate QTL underlying lung CFU. (A) Cross schema between CC030 and CC029 CC strains that contain CAST informative haplotypes at Chr 7 or 15. CC030 contains CAST at 30-45Mb on Chr 7 and CC029 contains CAST at 77-79Mb on Chr 15. The F2 population based on these founders were genotyped and informative segregants were infected with Mtb. (B) Lung CFU at 1-month post-in fection. F2 mice are grouped by the four possible genotypes: CAST at both 7 and 15, CAST at 7, CAST at 15, or non-CAST at both loci. To deter mine the effect of CAST haplotype on lung CFU, data was analyzed by 2-factor ANOVA to determine statistical significance.

### *Mtb* adapts to diverse hosts by utilizing distinct gene repertoires

This survey of disease-associated traits indicated that the CC panel encompasses a number of qualitatively distinct immune phenotypes. To determine if different bacterial functions were necessary to adapt to these conditions, we used TnSeq to estimate the relative abundance of individual *Mtb* mutants after selection in each CC host genotype. To serve as benchmarks of known immunological lesions, we also performed TnSeq in B6 mice that were lacking the mediators of Th1 immunity, lymphocytes (*Rag2*^-/-^) and IFNγ (*Ifnγ*^-/-^), or were lacking the immunoregulatory mediators that control disease by inhibiting inflammation, nitric oxide synthase (*Nos2*^-/-^) (Mishra et al., 2013) or the NADPH phagocyte oxidase (*Cybb*^-/-^) (Olive et al., 2018). The relative representation of each *Mtb* mutant in the input pool versus the pools recovered from mouse spleens was quantified (**Table S4**). Consistent with our previous work (Bellerose et al., 2020; Sassetti and Rubin, 2003), we identified 234 *Mtb* genes that are required for growth or survival in *Mtb* in B6 mice, based on significant underrepresentation of the corresponding mutant after four weeks of *in vivo* selection. All but one of these genes were found to be required in the larger panel, increasing confidence in this *Mtb* gene set (**Figure 4A** and **4B**). While the number of genes found to be necessary in each genotype across the panel was largely similar, the composition of these gene sets varied considerably. As more CC strains, and presumably more distinct immune states, were included in the analysis, the cumulative number of genes necessary for growth in these animals also increased. This cumulative gene set plateaued at ∼750, after the inclusion of approximately 20-25 genotypes (**Figure 4A**). This number of genes far outnumbered those identified in the panel of immunodeficient KO strains (**Figure 4B** and **Table S4**).

**Figure 4.**
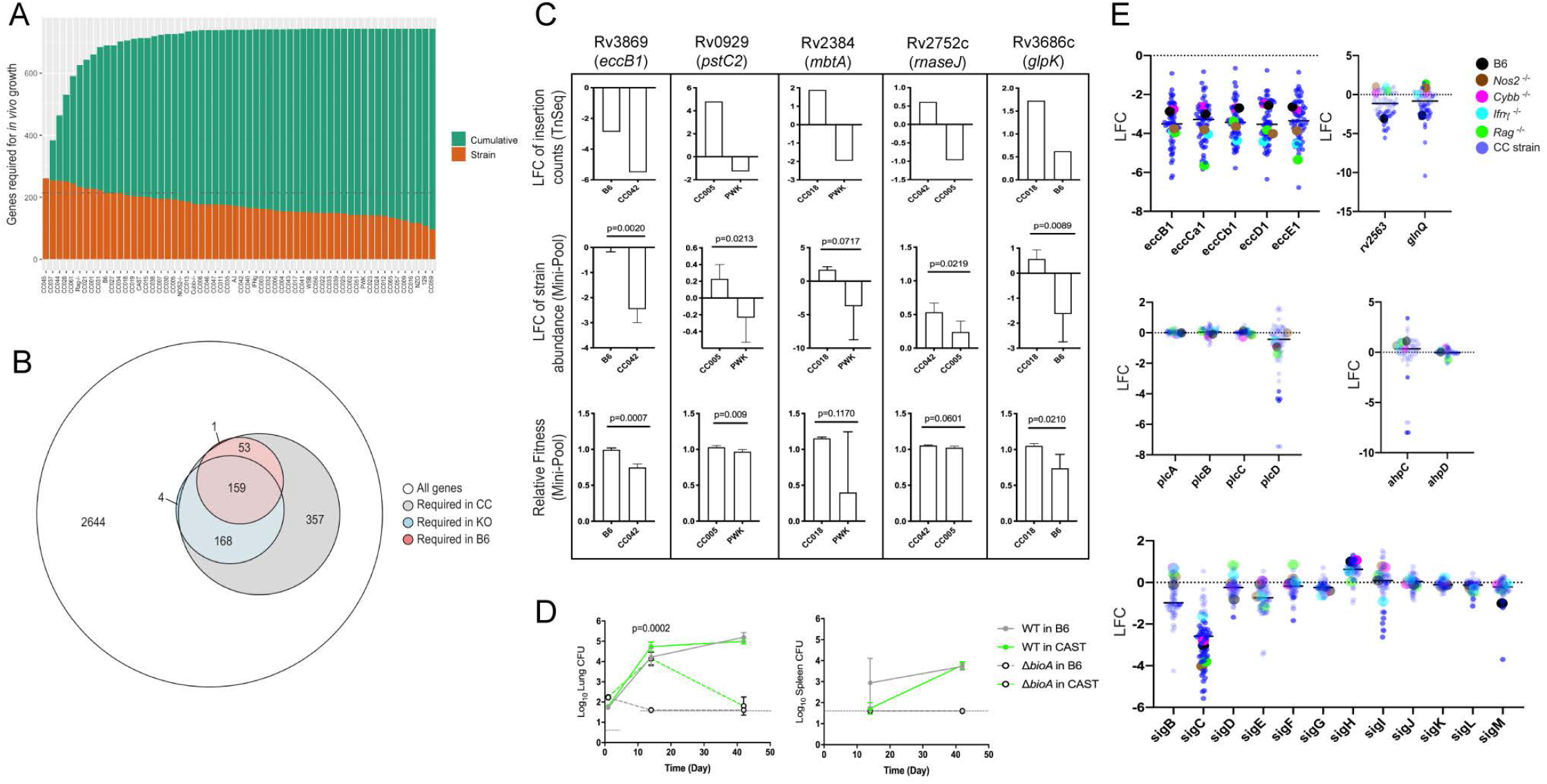
Mtb genetic requirements vary across diverse hosts. (A) The number of Mtb genes required for growth or survival in each diverse mouse strain across the panel (Qval ≤ 0 .05). Orange indicates the mutants required for each strain; turquoise shows the cumulative requirement as each new host strain is added. (B) Venn diagram showing the composition of Mtb gene sets required in each category of host (white, largest circle), only required in the CC panel (grey), required in specific immunological KO mice (blue) and genes required in B6 mice (red). In order to be called “essential” in each mouse strain, Mtb genes had to be significantly over or unrepresented in at least 2 genotypes. (C) Each box shows Log2 fold change (LFC) of individual mutants from the TnSeq screen relative to the input pool in indicated mouse strains (top); Log2 fold change of the indicated deletion mutants relative to WT from a pooled mutant validation infection (middle panel); Relative fitness calculated from (middle panel) to account for generation differences in each host due to differential growth rate. Bars are the average of 3-6 mice per mutant/genotype +/- SD. Statistical differences between mini-pool validation groups was assessed by Welch’s t test. (D) Lung CFU and spleen CFU from single strain low-dose aerosol infections of ∆bioA mutant or WT H37Rv strain in B6 and CAST mice at 2- or 5-weeks post-infection. Dashed line indicates the limit of detection. Each point indicates the average CFU +/- SD of 4-5 mice per group. (E) Log2 fold change of selected mutants from the TnSeq screen across the CC panel and immunological KO mice. Each dot represents the average LFC per mouse genotype. All mice in the complete mouse panel TnSeq screen were male; mice in the ∆bioA aerosol validation were female; mice in the mini-pool validation studies were male and female with no significant differences detected.

To verify that our TnSeq study accurately predicted the effect of the corresponding loss-of-function alleles, we assessed the phenotypes of selected bacterial deletion mutants in a small set of mouse genotypes that were predicted to result in differential selection. Individual *Mtb* mutants lacking genes necessary for ESX-1 type VII secretion (*eccB1*), siderophore-mediated iron acquisition (*mbtA*), phosphate transport (*pstC2*), glycerol catabolism (*glpK*), and RNA processing (*rnaseJ*) were generated and tagged with a unique molecular barcode. These mutants were combined with a barcoded wild-type parental strain, the resulting ‘mini-pool’ was subjected to *in vivo* selection in the same manner as the TnSeq study, and the relative abundance of each mutant was determined by sequencing the amplified barcodes. In each case, the difference in relative abundance predicted by TnSeq was reproduced with deletion mutants (**Figure 4C**). In this simplified system, we were able to accurately quantify the expansion of the bacterial population and calculate the “fitness” of each mutant relative to the wild-type strain. Fitness reflects the inferred doubling time of the mutant, where a fitness of 1 is defined as wild-type, and 0 represents a complete lack of growth. Even by this metric, the deletion mutants displayed the differences in fitness between mouse strains that was predicted by TnSeq (**Figure 4C**). The statistical significance of these differences in abundance or fitness were similar for each mutant (between p=0.009 and p=0.06), except for *mbtA* where the variation was higher, and confidence was modestly lower (p=0.07 and p=0.12). This study also allowed us to estimate the sensitivity of the TnSeq method, which could detect even the 30% fitness defect of the ∆*glpK* strain between the B6 and CC018 animals (**Figure 4C**), a defect that was not observed in previous studies in BALB/c mice (Bellerose et al., 2019; Pethe et al., 2010).

To also validate TnSeq predictions in a single-strain aerosol infection model, we used a biotin biosynthetic mutant. *bioA* is necessary for biotin production and is essential for growth in B6 mice (Woong Park et al., 2011). Our TnSeq study (**Table S4**) predicted this mutant was less attenuated in the CAST background (ratio of input/selected = 12.1) than in the B6 strain (ratio of input/selected = 42.2). Two weeks after aerosol infection, we found that the ∆*bioA* mutant was cleared from the lungs and spleen of B6 mice but displayed similar growth to wild-type in the lungs of CAST mice (**Figure 4D**). By 6 weeks post infection the ∆*bioA* mutant had also been largely cleared from the lungs of CAST (**Figure 4D**). Thus, while TnSeq was unable elucidate the complex kinetics of this phenotype, it accurately predicted the relative levels of growth attenuation in these host backgrounds.

### The immunological diversity of CC mice is reflected in the pathogen’s genetic requirements

The distribution of *Mtb*’s requirements across the mouse panel suggested the presence of two broad categories of genes. A set of 136 “core” virulence functions were required in the majority of mouse genotypes, and a second larger set of 607 “adaptive” virulence genes were required in only a subset of lines (**Table S4**). The core functions included a number of genes previously found to be important in B6 mice, including those necessary for the synthesis of essential cofactors, such as pyridoxine (*Pdx1*) (Dick et al., 2010); for the acquisition of nutrients, such as siderophore-bound iron (*irtAB*) (Ryndak et al., 2010), cholesterol (*mce4*) (Pandey and Sassetti, 2008), glutamine (*glnQ* and *rv2563*) (Bellerose et al., 2020); and for type VII secretion (ESX1 genes) (Stanley et al., 2003). Despite the importance of these core functions, a large range in the relative abundance of these mutants was observed across the panel, and in some cases specific immunological requirements could be discerned. Mutants lacking the major structural components of the ESX1 system were attenuated for growth in B6 mice, as expected. This requirement was consistently enhanced in mice lacking *Rag2, Ifnγ*, or *Nos2* (**Figure 4E**), consistent with the preferential role of ESX1 during the initial stage of infection before the initiation of adaptive immunity (Stanley et al., 2003). In contrast, the attenuation of mutants lacking the *glnQ* encoded glutamine uptake system was relieved in all four immunodeficient mouse lines (**Figure 4E**). In both cases, the differential mutant abundance observed in these KO mice was reproduced, or exceeded, in the CC panel.

The adaptive virulence functions included a number of genes previously thought to be dispensable in the mouse model and were only necessary in CC strains. For example, the alkyl hydroperoxide reductase, AhpC has been proposed to function with the adjacently encoded peroxiredoxin, AhpD and is critical for detoxifying reactive nitrogen intermediates *in vitro* (Chen et al., 1998; Hillas et al., 2000). However, deletion of *ahpC* has no effect on *Mtb* replication in B6 or BALB/c mice (Springer et al., 2001), and we confirmed that *ahpC* and *ahpD* mutations had no effect in any of the B6-derived lines. In contrast, *ahpC*, but not *ahpD* mutants were highly attenuated in a small number of CC strains (**Figure 4E**). Similarly, the four phospholipase C enzymes of *Mtb* (*plcA-D*) are implicated in both fatty acid uptake and modifying host cell membranes but are dispensable for replication in B6 mice (Le Chevalier et al., 2015). Again, while we found that none of these genes were required in B6-derived KO lines, the *plcD* mutants were specifically underrepresented in a number of CC mice (**Figure 4E**). These individual bacterial functions are controlled by regulatory proteins, such as the extracytoplasmic sigma factors. Despite the importance of these transcription factors in the response to stress, only *sigF* has been consistently been shown to contribute to bacterial replication in standard inbred lines of mice (Geiman et al., 2004; Rodrigue et al., 2006). Our study assesses the importance of each sigma factor in parallel across diverse host genotypes and identified a clear role for several of these regulators. *sigC, sigI, sigF, sigL*, and *sigM* mutants were each significantly underrepresented in multiple lines of mice, and several of these phenotypes were only apparent in the diverse CC animals (**Figure 4E**). In sum, the 607 adaptive functions that are differentially required across the host panel represents nearly 20% of the non-essential gene set of Mtb, suggesting that a significant fraction of the pathogen’s genome is dedicated to maintaining optimal fitness in diverse host environments.

### Differential genetic requirements define virulence pathways in *Mtb*

To more formally investigate the distinct stresses imposed on the bacterial population across this host panel, we characterized the differentially required bacterial pathways. Upon performing each possible pairwise comparison between the *in vivo* selected mutant pools, we found 679 mutants whose representation varied significantly (FDR < 5%) in at least two independent comparisons (**Table S4**). We then applied weighted gene correlation network analysis (WGCNA) (Langfelder and Horvath, 2008) to divide the mutants into 20 internally-correlated modules. Further enrichment of these modules for the most representative genes (intramodular connectivity > 0.6) revealed that nearly all modules contained genes that are encoded adjacently in the genome and many of these modules consisted of genes dedicated to a single virulence-associated function (**Figure 5A**). Module 3 contains two distally encoded loci both known to be necessary for ESX1-mediated protein secretion, the primary ESX1 locus (*rv3868-rv3883*) and the *espACD* operon (*rv3616c-rv3614c*). Similarly, other modules consisted of genes responsible for ESX5 secretion (Module 7), mycobactin synthesis (Module 4), the Mce1 and Mce4 lipid importers (Modules 5 and 16), phthiocerol dimycocerosate synthesis (PDIM, Module 8), PDIM transport (Module 16), and phosphate uptake (Module 14).

**Figure 5.**
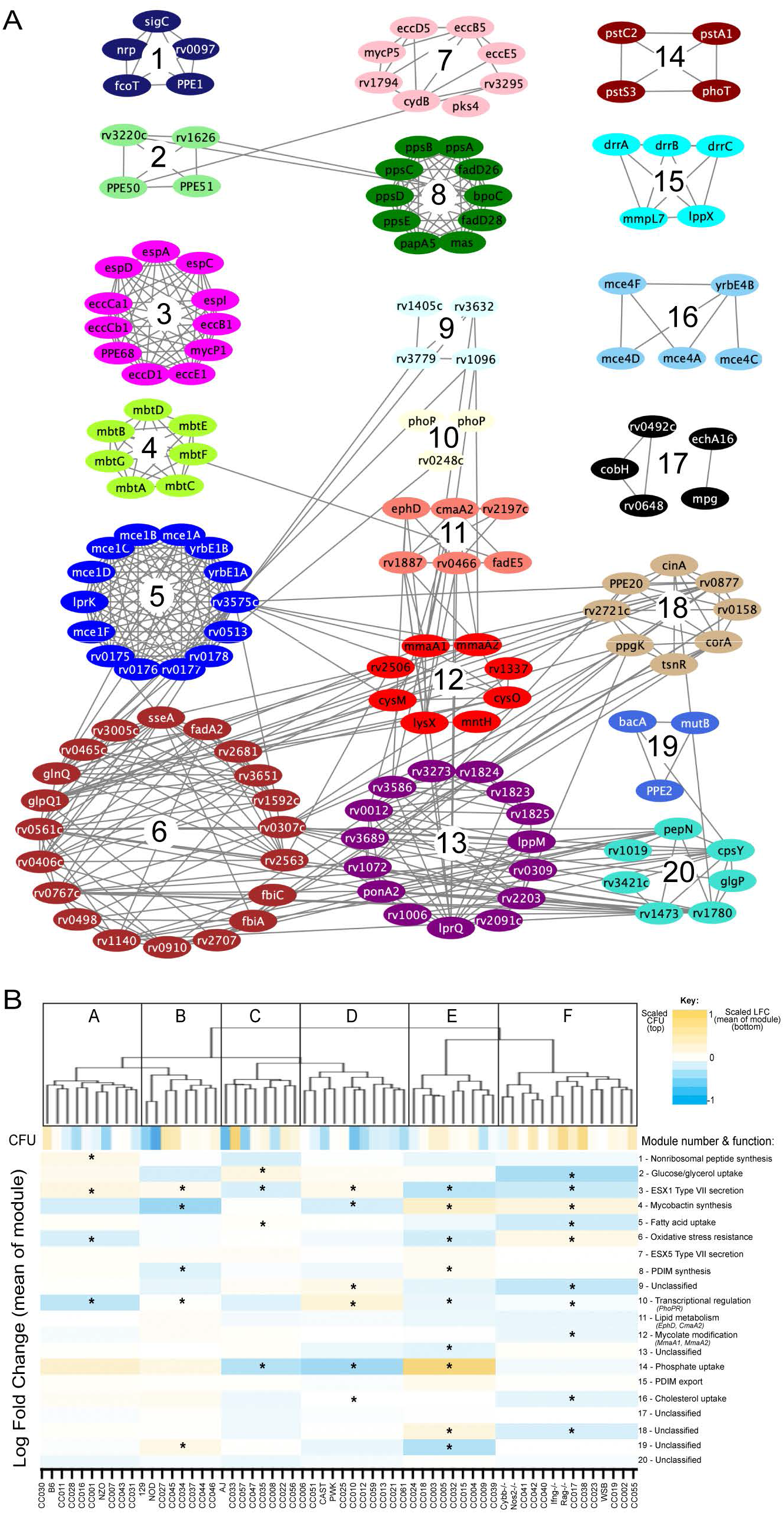
Mtb virulence pathways associate with distinct host immune pressures. (A) Weighted gene correlation network analysis (WGCNA) of the 679 Mtb genes that significantly vary across the diverse mouse panel. The most representative genes of each module (intramodular connectivity > 0.6) are shown. (B) Mouse genotypes were clustered based on the relative abundance (log fold change relative to in vitro) of the 679 variable Mtb mutants. The six major clusters (Cluster A-F) were associated with both CFU and the relative abundance of mutants in each bacterial module (1-20; right hand-side with known functions). Cluster F is associated with high CFU (p<0.05, as described in methods). ∊ indicate modules significantly associated with specific mouse clusters (P<0.05).

The 20 genes assigned to Module 6 included two components of an important oxidative stress resistance complex (*sseA* and *rv3005c*) and were highly enriched for mutants predicted to be involved in this same process via genetic interaction mapping (11/20 genes were identified in (Nambi et al., 2015), a statistically significant overlap (p< 2.8e-10 by hypergeometric test)). Thus, each module represented a distinct biological function.

Many pathway-specific modules contained genes that represented novel functional associations. For example, the gene encoding the sigma factor, *sigC*, was found in Module 1 along with a non-ribosomal peptide synthetic operon. Previous genome-wide ChIP-seq and overexpression screens support a role for SigC in regulating this operon (Minch et al., 2015; Turkarslan et al., 2015). Similarly, *rv3220c* and *rv1626* have been proposed to comprise an unusual two component system that is encoded in different regions of the genome (Morth et al., 2005). Both of these genes are found in Module 2, along with the PPE50 and PPE51 genes that encode at least one outer membrane channel (Wang et al., 2020)(**Figure 5A**). In both cases, these associations support both regulatory and obligate functional relationships between these genes. Six of the 20 modules were not obviously enriched for genes of a known pathway, demonstrating that novel virulence pathways are important for adapting to changing host environments.

To explore the complexity of immune environments in the CC, we used the TnSeq profiles of the 679 differentially fit *Mtb* mutants to cluster the mouse panel into 6 major groups of host genotypes (**Figure 5B**). One mouse cluster was significantly associated with high CFU (Cluster F, **Figure 5B**), which contained susceptible *Nos2*^-/-^, *Cybb*^-/-^, *Ifnγ* ^-/-^, and *Rag2*^-/-^ animals. This high CFU cluster was associated with alterations in a diverse set of bacterial modules and corresponded to an increased requirement for lipid uptake (Modules 5 and 16) and ESX1, consistent with previous TnSeq studies in susceptible *Nos2*^-/-^ and C3HeB/FeJ mice (Mishra et al., 2017). In addition, we identified a significant reduction in the requirement for the oxidative stress resistance (Module 6) in the highest CFU cluster. Despite these associations between bacterial genetic requirements and susceptibility, the clustering of mouse genotypes was largely independent of overall susceptibility. Similarly, while Module 1 was significantly associated with high IFNγ levels, other bacterial fitness traits were not highly correlated with cytokine abundance (**Figure S3)**. Instead, each major mouse cluster was associated with a distinct profile of *Mtb* genetic requirements. This observation supported the presence of qualitatively distinct disease states and complex genetic control of immunity.

### Identification of host-pathogen genetic interactions

To investigate the host genetic determinants of the bacterial microenvironment, we associated *Mtb* mutant fitness profiles with variants in the mouse genome. When the relative abundance of each *Mtb* mutant was considered individually, the corresponding “Host Interacting with Pathogen QTL” (*Hip*QTL) were distributed across the mouse genome (**Figure 6A**). 41 of these traits reached an unadjusted p-value threshold of 0.05 and can be considered as robust for single hypothesis testing (*Hip1*-*41*, **Table S2** and **S5**). These included *Hip*QTL associated with both *ahpC* and *eccD1*, that explain at least a portion of the observed variable abundance of these mutants (**Figure 4E**). In order to reduce complexity and increase the power of this analysis, we performed QTL mapping based on the first principal component of each of the previously defined modules of *Mtb* virulence pathways (**Figure 5A**). Three of these “eigentraits” were associated with QTL at a similar position on chromosome 10 (**Figure 6B**), corresponding to Module 3 (TypeVII secretion, ESX1), Module 4 (mycobactin synthesis, *mbt*), or Module 16 (cholesterol uptake, *mce4*). In all three cases, a single mutant from the module was independently associated with a QTL at the same position as the module eigentrait (**Table S5**, *Hip21, Hip22, Hip24*), and all genes in the corresponding network cluster (**Figure 5A**) mapped to the same location (**Figure 6C**-**E**). While not all individual traits mapped with high confidence, the coincidence of these multiple QTL was statistically significant (**Figure 6B**).

**Figure 6.**
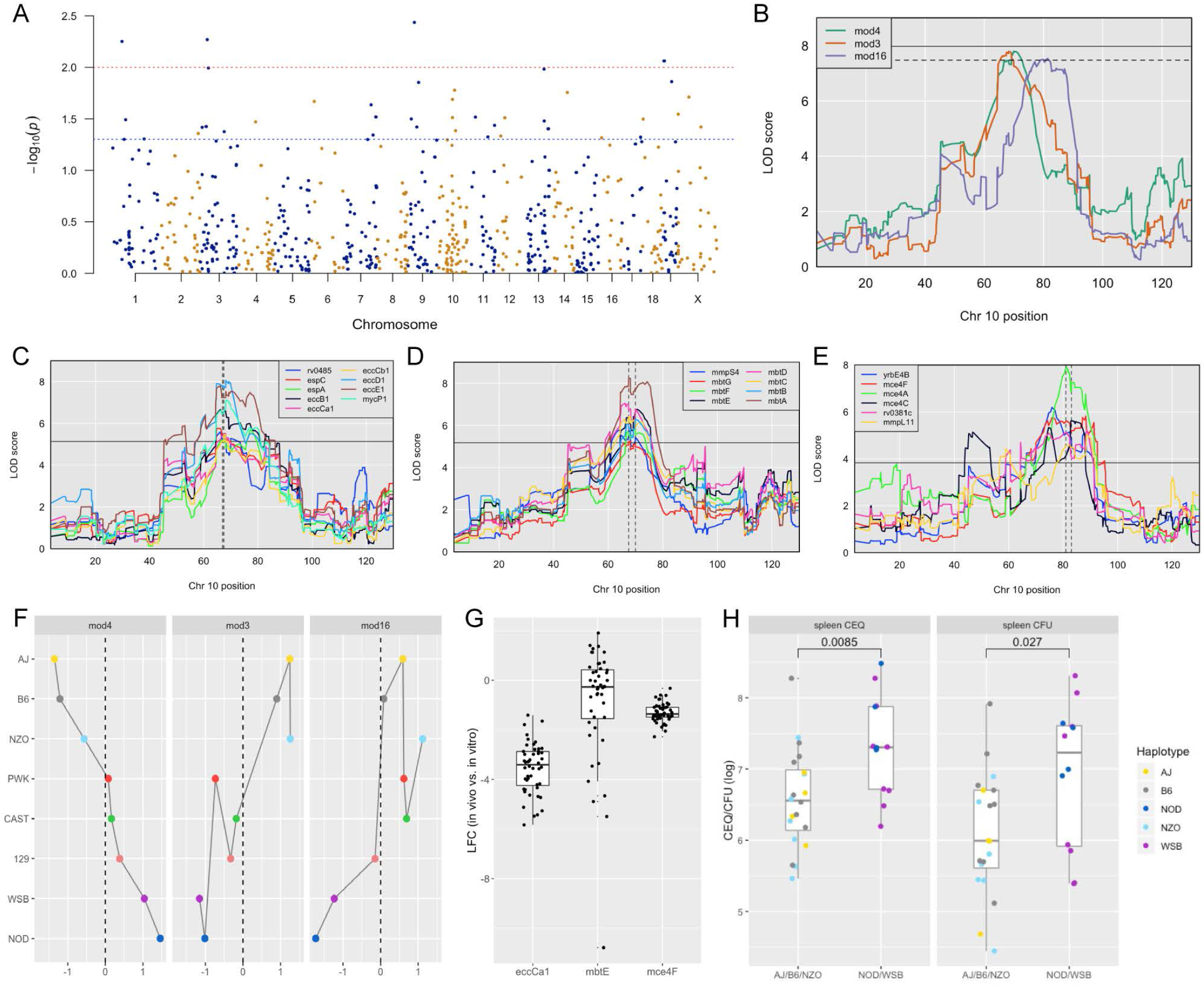
Identification of ‘Host Interacting with Pathogen’ QTL mapping (HipQTL). (A) Manhattan plot of single Mtb mutant QTL mapping across the mouse genome. Each dot represents an individual Mtb mutant plotted at the chromosomal location of its maximum LOO score. Red dashed line indicates P<0.01; Blue P<0.05. (B) Chromosome 10 QTL corresponding to Mtb eigentraits identified in network analysis in Figure 5. Module 3 (type VII secretion, ESX1 operon; orange), Module 4 (Mycobactin synthesis, mbt; green) and Module 16 (Cholesterol uptake, mce4; purple) are shown. Solid and dotted lines indicated P = 0.05 and P = 0.1, respectively. (C-E) QTL mapping of single Mtb mutants corresponding to the (C) ESX1 module, (D) mbt module and (E) mce4 modules. Coincidence of multiple QTL was assessed by the NL-method of Neto et al (Neto et al., 2012). Thresholds shown are for N=9, N=8, and N=6 for panels C, D, and E, respectively. (F) Parental founder effects underlying Module 3, 4 and 16 QTL. Allele effects were calculated at the peak LOD score marker on chromosome 10. (G) Distribution of log fold change (LFC) of representative single mutants from each module; eccCa1 (ESX1 module), mbtE (mbt module) and mce4F (mce4 module) relative to in vitro. Each dot is the LFC of the specified mutant in each CC mouse strain. Box and whiskers plots (Tukey) of each trait indicate the median and interquartile range. (H) Spleen CEQ and Spleen CFU for CC strains (box plots as in G). Mouse values are grouped by the parental haplotype allele series underlying the chromosome 10 Hip42 locus (NOD/WSB vs AJ/B6/NZO). Each dot represents the average CFU/CEQ of each CC genotype. Statistical differences in disease-associated traits and distinct haplotypes groups was assessed by t-test. LOD, logarithm of the odds; LFC, log fold change; CEQ, Chromosomal equivalents; CFU, colony forming units.

Both the relative positions of the module-associated QTL and the associated founder haplotypes indicated that a single genetic variant controlled the abundance of ESX1 and *mbt* mutants (*Hip42*). Specifically, we found no statistical support for differentiating these QTL based on position (p=0.93)(Boehm et al., 2019) and the same founder haplotypes were associated with extreme trait values at both loci, though they had opposite effects on the abundance of ESX1 mutants and *mbt* mutants (**Figure 6F**). We conclude that a single haplotype has a pleiotropic effect on *Mtb*’s environment and has opposing effects of the requirement for mycobactin synthesis and ESX1 secretion. The relationship between this variant and the *mce4*-associated QTL (*Hip43*) was less clear, as the statistical support for independent QTL was weak (ESX1 and *mce4* QTL p=0.17; *mbt* and *mce4* QTL p=0.08) and the effects of founder haplotypes were similar but not identical (**Figure 6F**). Some of this ambiguity may be related to the relatively small range in trait values for *mce4*, compared to either ESX1 or *mbt* (**Figure 6G**). Based on this data, we report two distinct *Hip*QTL in this region (*Hip42* and *43*, **Table S5**).

Two *Tip*QTL overlapped with *Hip*QTL (**Figure S2**), suggesting specific interactions between bacterial fitness and immunity. However, most *Tip*- and *HipQTL* were distinct, indicating that the fitness of sensitized bacterial mutants can be used to detect genetic variants that subtly influence the bacterial environment but not overtly alter disease. We chose to further investigate whether *Hip*QTL might alter overall bacterial disease using the most significant *Hip*QTL on chromosome 10 (*Hip42*). We found that the founder haplotypes associated with extreme trait values at this QTL could differentiate CC strains with significantly altered total bacterial burden, and the NOD and WSB haplotypes were associated with higher bacterial numbers (p=0.0085 for spleen CEQ; p=0.027 for spleen CFU; **Figure 6H**). Thus, not only could the *Hip*QTL strategy identify specific interactions between host and bacterial genetic variants, but it also appears to be a sensitive approach for identifying host loci that influence the trajectory of disease.

### Identifying candidate genes underlying QTL

A pipeline was designed to prioritize genetic variants based on genomic and tuberculosis disease criteria. We concentrated on three QTL: two that were highly significant and with clear allele effects (*Tip5, Hip42*), and the *Tip8* locus which we validated by intercross. For each QTL region, we identified genes that belonged to a differentially expressed transcriptional module in mouse lungs following *Mtb* infection (Moreira-Teixeira et al., 2020). Next, we identified genetic variants segregating between the causal CC haplotypes in the gene bodies corresponding to these transcripts, and prioritized missense or nonsense variants.

For the *Tip5* QTL underlying CEQ, CFU, and IL-10 levels, we identified nine candidate genes with regulatory or splicing variants and two genes with missense variants specific to the NOD haplotype. Of these candidates, cathepsin Z (*Ctsz)* encodes a lysosomal cysteine proteinase and has previously been associated with TB disease risk in humans (Adams et al., 2011; Cooke et al., 2008). The QTL underlying lung CFU and CXCL1 abundance (*Tip8*), which was driven solely by the genetically divergent CAST founder haplotype, contained over 50 genes (**Table S6**) and will need further refinement. The QTL associated with the abundance of ESX1 and *mbt* mutants (*Hip42*) had a complex causal haplotype pattern (AJ/B6/NZO vs. 129/CAST/PWK vs. NOD/WSB) suggesting multiple variants might be impacting common genes. Within this interval, we identified 13 genes expressed in response to *Mtb* infection, three of which had SNPs fully or partially consistent with at least one of the identified causal haplotype groups (**Table S6**). *Ank3* contains several SNPs in the 3’ UTR and other non-coding exons that differentiated NOD/WSB from the other haplotypes. Similarly, *Fam13c* had two missense mutations following the same haplotype pattern. For the AJ/B6 haplotype state, we identified a missense mutation and several variants in the 3’ UTR of *Rhobtb1*, which belongs to the Rho family of the Ras superfamily of small GTPases (Goitre et al., 2014). Overall, the evidence supports a role for *Rhobtb1* in a monogenic effect at the chromosome 10 locus. This evidence includes both protein coding differences dividing AJ/B6 from the other haplotypes, a potential expression/transcript regulatory difference that segregates the NOD/WSB state from the remaining parental haplotypes, and a plausible role for this gene in controlling intracellular trafficking (Long et al., 2020) and the opposing requirements for ESX1 and mycobactin.

## Discussion

Our immunological analysis of the CC panel identified correlates of TB disease progression that were consistent with previous studies in both mice and human patients (Ahmed et al., 2020; Niazi et al., 2015; Zak et al., 2016). More importantly, we identified outlier strains that produce distinct immunological states, highlighting how our previous reliance on genetically homogenous lab strains of mice has oversimplified our understanding of TB pathogenesis. For example, despite the generally strong correlation between lung bacterial burden and disease, CC030/GeniUnc and CC040/TauUnc mice suffered from more inflammation and wasting than would be predicted from the number of bacteria in their lungs or spleens. This phenotype reflects a failure of disease “tolerance”, which is proposed to be a critical determinant of protective immunity (Ayres and Schneider, 2012; Olive et al., 2018). Similarly, we identified a number of CC genotypes that produce very low, or undetectable, levels of the protective cytokine IFNγ, but still control lung bacterial replication. While a growing body of literature suggests that immune responses distinct from the canonical Th1 response can control infection (Lu et al., 2019; Sakai et al., 2016), these CC strains are the first example of an animal model in which IFNγ appears to be dispensable. The reproducibility of the CC strains facilitated the identification of these phenotypes and provides tractable models for their characterization.

The ability to separate aspects of the immune response from disease progression implied that these features are under distinct multigenic control. This conclusion is supported by genetic mapping, which identified a number of variants that control distinct aspects of the immune response to *Mtb*. The QTL identified in this study are generally distinct from CC loci that control immunity to viruses (Ferris et al., 2013; Gralinski et al., 2017; Noll et al., 2020) or another intracellular bacterial pathogen, *Salmonella* (Zhang et al., 2019). However, *Tip8* and *Tip10* overlap with QTL previously defined via *Mtb* infection of CC001xCC042 F_2_ intercross population (Smith et al., 2019) suggesting that common variants may have been identified in both studies. While the specific genetic variants responsible for these QTL remain unknown, both coincident trait mapping and bioinformatic analysis suggest mechanistic explanations for some QTL-phenotype associations. For example, a single interval on chromosome 2 controls CFU levels and IL-10, and contains a variant in the *Ctsz* gene encoding Cathepsin Z. *Ctsz* is a strong candidate considering its known roles in autophagy (Amaral et al., 2018), dendritic cell differentiation and function (Obermajer et al., 2008), its upregulation in non-human primates (Ahmed et al., 2020) and human patients with *Mtb* (Zak et al., 2016), and the association of *CTSZ* variants with disease risk in human TB studies (Adams et al., 2011; Cooke et al., 2008).

Using TnSeq as a multidimensional phenotyping method across this population provided insight into how the diversity of host-derived microenvironments have shaped the pathogen’s genome. While *Mtb* is an obligate pathogen with no significant environmental niche, only a minority of the genes in its genome have been found to contribute to bacterial fitness in either laboratory media or individual inbred mouse models, leaving the pressures that maintain the remaining genomic content unclear. We find that approximately three times more genes contribute to bacterial growth or survival in the CC population than in the standard B6 model. While some bacterial genetic requirements could be associated with known immune pathways, most of the differential pressures on bacterial mutants could not be attributed to these simple deficiencies in known mechanisms of immune control. Instead, it appears that the CC population produces a spectrum of novel environments, and that a relatively large fraction of the pathogen’s genome is needed to adapt to changing immune pressures. Differential pressures on these adaptive virulence functions are similarly apparent in genomic analyses of *Mtb* clinical isolates. Signatures of selection have been detected in ESX1-related genes (Holt et al., 2018; Sousa et al., 2020), *phoPR* (Gonzalo-Asensio et al., 2014), and the oxidative stress resistance gene *sseA* (de Keijzer et al., 2014), suggesting that *Mtb* is exposed to similarly variable host pressures in genetically diverse human and mouse populations. While the combinatorial complexity of associating host and pathogen genetic variants in natural populations is daunting, the identification of *Hip*QTL in the CC panel indicates that these inter-species genetic interactions can be important determinants of pathogenesis and can be dissected using tractable models of diversity.

## Acknowledgements

We thank all members of Sassetti Lab, past and present for technical help and discussions; Dr. Nathan Hicks and Dr. Sarah Fortune for kindly providing the *rnaseJ* mutant; Dr. David Tobin for insightful manuscript comments; Dr. Dennis Ko for QTL acronym creativity; Erin Curtis for mouse schema expertise and the Systems Genetic Core at UNC for their help in procuring CC mice in timely fashion. This work was supported by NIH grants AI132130 to C. M. Sassetti and FPMV; U19AI100625 to FPMV and MTF; a fellowship from the Charles H. King Foundation to C. M. Smith; and a HHMI Gilliam Fellowship A20-0146 to BKH. The genetic characterization of the CC strains was supported in part by NIH grant U24HG010100 to FPMV.

## Author contributions

Conceptualization, C.M. Smith and C.M Sassetti; Methodology, R.E.B, K.C.M, K.P, T.R.L; Investigation, C.M. Smith, M.K.P, B.B.M, J.E.L, M.C.K, M.M.B, A.J.O, C.J.R, C.M. Sassetti; Validation, S.W.P, H.L, S.E, D.S; Formal Analysis, R.E.B, F.J.B, R.K.M, B.K.H, C.L, M.T.F, T.R.I; Resources, G.D.S, P.H, T.A.B; Visualization, C.M. Smith, R.E.B, M.K.P, R.K.M, C.M. Sassetti; Writing – original draft, C.M Sassetti and C.M Smith; Writing – review and editing, all authors; Supervision, C.M. Sassetti; Funding acquisition, C.M. Sassetti, M.T.F, F.P.M.V.

## Materials and Methods

### Ethics statement

Mouse studies were performed in strict accordance using the recommendations from the Guide for the Care and Use of Laboratory Animals of the National Institute of Health and the Office of Laboratory Animal Welfare. Mouse studies at the University of Massachusetts Medical School (UMASS) were performed using protocols approved by the UMASS Institutional Animal Care and Use Committee (IACUC) (Animal Welfare Assurance Number A3306-01) in a manner designed to minimize pain and suffering in *Mtb*-infected animals. Any animal that exhibited severe disease signs was immediately euthanized in accordance with IACUC approved endpoints. All mouse studies at UNC (Animal Welfare Assurance #A3410-01) were performed using protocols approved by the UNC Institutional Animal Care and Use Committee (IACUC).

### Mice

Male and female Collaborative Cross parental strains (A/J #0646; C57BL/6J #0664; 129S1/SvImJ #02448; NOD/ShiLtJ #01976; NZO/HiLtJ #02105; CAST/EiJ #0928, PWK/PhJ #3715 and WSB/EiJ #01145) and single gene immunological knockout mice were purchased from The Jackson Laboratory (*Nos2*^-/-^ #2609, *Cybb*^-/-^ #2365, *Ifnγ*^-/-^ #2287) or Taconic (*Rag*N12) and bred at UMASS. Male mice from 52 CC strains were purchased from the UNC Systems Genetics Core Facility (SGCF) between July 2013 and August 2014. The 52 CC strains used in this study include: CC001/Unc, CC002/Unc, CC003/Unc, CC004/TauUnc, CC005/TauUnc, CC006/TauUnc, CC007/Unc, CC008/GeniUnc, CC009/Unc, CC010/GeniUnc, CC011/Unc, CC012/GeniUnc, CC013/GeniUnc, CC015/Unc, CC016/GeniUnc, CC017/Unc, CC018/Unc, CC019/TauUnc, CC021/Unc, CC022/GeniUnc, CC023/GeniUnc, CC024/GeniUnc, CC025/GeniUnc, CC027/GeniUnc, CC028/GeniUnc, CC029/Unc, CC030/GeniUnc, CC031/GeniUnc, CC032/GeniUnc, CC033/GeniUnc, CC034/Unc, CC035/Unc, CC037/TauUnc, CC038/GeniUnc, CC039/Unc, CC040/TauUnc, CC041/TauUnc, CC042/GeniUnc, CC043/GeniUnc, CC044/Unc, CC045/GeniUnc, CC046/Unc, CC047/Unc, CC051/TauUnc, CC055/TauUnc, CC056/GeniUnc, CC057/Unc, CC059/TauUnc, CC060/Unc, CC061/GeniUnc, CC065/Unc, CC068/TauUnc. More information regarding the CC strains can be found at http://csbio.unc.edu/CCstatus/index.py?run=AvailableLines.information.

CC030 x CC029 F_2_ mice were generated at UNC by crossing CC030 and CC029 mice (purchased from the SGCF in 2016) to generate F_1_ s (both CC030 dam by CC029 sires as well as CC029 dam by CC030 sires). The resulting F_1_ s were subsequently intercrossed to generate F_2_ mice with all possible grandparental combinations. Male and female F_2_ mice were shipped to UMASS for *Mtb* infections.

All mice were housed in a specific pathogen-free facility under standard conditions (12hr light/dark, food and water ad libitum). Mice were infected with *Mtb* between 8-12 weeks of age. Male and female mice were used, unless otherwise noted.

### M.tuberculosis strains

All *M*. *tuberculosis* strains were grown in Middlebrook 7H9 medium containing oleic acid-albumin-dextrose-catalase (OADC), 0.2% glycerol, and 0.05% Tween 80 to log-phase with shaking (200 rpm) at 37°C. Hygromycin (50 ug/ml) or kanamycin (20 ug/ml) was added when necessary. The ∆*bioA* strain was made by homologous recombination as previously described (Woong Park et al., 2011). For pooled mutant infections, deletion strains were constructed using ORBIT (Murphy et al., 2018), which included gene replacement by the vector pKM464 carrying unique q-Tag sequences to identify each mutant for deep sequencing. The *rnaseJ* mutant was also made by ORBIT and was kindly provided by Dr. Nathan Hicks and Dr Sarah Fortune. Prior to all *in vivo* infections, cultures were washed, resuspended in phosphate-buffered saline (PBS) containing 0.05% Tween 80, and sonicated before diluting to desired concentration (see below).

### Mouse Infections

For TnSeq experiments, 1×10^6^ CFU of saturated *Himar1* transposon mutants (Sassetti et al., 2003) was delivered via intravenous tail vein injection. Mice were infected over 3 infection batches, as denoted in **Table S1**). At indicated time points mice were euthanized, and organs were harvested then homogenized in a FastPrep-24 (MP Biomedicals). CFU was determined by dilution plating on 7H10 agar with 20 ug/mL kanamycin. For library recovery, approximately 1×10^6^ CFU per mouse was plated on 7H10 agar with 20ug/mL kanamycin. After three weeks of growth, colonies were harvested by scraping and genomic DNA was extracted. The relative abundance of each transposon mutant was estimated as described (Long et al., 2015).

Single strain aerosol infections were performed in a Glas-Col machine to deliver 50-150 CFU/mouse. At indicated time points, mice were euthanized, and organs were harvested then homogenized in a FastPrep-24 (MP Biomedicals). CFU was determined by dilution plating on 7H10 agar with 20 ug/mL kanamycin or 50ug/mL hygromycin as required.

Chromosomal equivalent (CEQ) was enumerated according to previously published protocol (Lin et al., 2014; Muñoz-Elías et al., 2005). Cytokines and chemokines were assayed from organ homogenates using the pro-inflammatory focused 32-plex (Eve Technologies, Calgary, CA) or IFNγ DuoSet ELISA (R&D Systems) according to manufacturer protocol. IFNγ release was assayed by ELISPOT (BD Biosciences #551083) according to manufacturer protocol. Single cell suspensions of lung or spleen tissue were stimulated for 18-24 hours with 4ug/ml purified protein derivative (PPD) and after development, spots were quantified using a CTL Immunospot S5 Analyzer.

For pooled mutant infections, mice were infected with a pool of deletion mutants at equal ratios via the intravenous route (1×10^6^ CFU/mouse). At indicated time points, approximately 10,000 CFU from the spleen homogenate of each mouse was plated on 7H10 agar. Genomic DNA was extracted for sequencing as described previously (Long et al., 2015). Sequencing libraries spanning the variable region of each q-Tag were generated using PCR primers binding to regions common among all q-Tags, similar to previously described protocols (Bellerose et al., 2020; Martin et al., 2017). In each PCR, a unique molecular counter was incorporated into the sequence to allow for the accurate counting of input templates and account for PCR jackpotting. The libraries were sequenced to 1,000-fold coverage on an Illumina NextSeq platform using a 150-cycle Mid-Output kit with single-end reads. Total abundance of each mutant in the library was determined by counting the number of reads for each q-Tag with a unique molecular counter. Relative abundance of each mutant in the pool was then calculated by dividing the total abundance of a mutant by the total abundance of reads for wild-type H37Rv. The relative abundance was normalized to relative abundance at initial infection (Day 0) and log_2_ transformed. Fitness was calculated as previously described (Palace et al., 2014).

## QUANTIFICATION AND STATISTICAL ANALYSIS

### TnSeq analysis

TnSeq libraries were prepared and counts of each transposon mutant were estimated as described (Long et al., 2015). NCBI Reference Sequence NC_018143.1 was used for H37Rv genome and annotations. In the majority of cases, two replicate mouse libraries were used per genotype. Only a single TnSeq library was obtained for CC010, CC031, CC037, CC059, CC016, and PWK/PhJ. Insertion mutant counts across all libraries were normalized by beta-geometric correction (DeJesus et al., 2015), binned by gene, and replicate values for each mouse genotype averaged. Mean values for each gene were divided by the grand mean then log_2_ transformed and quantile normalized. The resulting phenotype values were used for both WGCNA and QTL mapping.

To eliminate genes having no meaningful variation across the mouse panel, statistical tests of log_2_ fold change (LFC) in counts between all possible pairs of mouse genotypes were performed by resampling (DeJesus et al., 2015). 679 “significantly varying genes” were identified whose representation varied significantly (FDR < 5%) in at least two independent comparisons. For relative mutant abundance estimates, LFC in counts between *in vitro*-grown H37Rv (6 replicate libraries) vs libraries from each mouse genotype were determined by resampling as above.

### WGCNA analysis

Weighted gene correlation network analysis (WGCNA) was applied to categorize the 679 significantly varying genes into 20 internally-correlated modules (Langfelder and Horvath, 2008). Modules were filtered (intramodular connectivity > 0.6) to obtain the most representative genes. First principal component scores of module eigengenes were used as phenotype values for QTL mapping after first winsorizing (q=0.05) using the R package *broman* (https://cran.r-project.org/web/packages/broman/index.html).

In order to perform association analysis between modules of genes and clusters of mice (**Figure 5B**), the mice were clustered based on the matrix of TnSeq LFCs for significantly varying genes using *hclust* in R (with the “Ward.D2” distance metric). Then, for each module of genes, the LFCs in each cluster of mice were pooled and compared to all the other mice using a T-test, identifying modules with a mean LFC in a specific mouse cluster that is significantly higher or lower than the average across all the other mice. The resulting p-values over all combinations of gene modules and mouse clusters were adjusted using Benjamini-Hochberg for an overall FDR < 0.05.

### Disease-related trait analysis and Heritability estimation

For the trait heatmap, trait values were clustered (*hclust* in R package *heatmaply;* traits scaled as per default function) and dendrogram nodes colored by 3 k-means. Correlation between disease-related TB traits was determined by Pearson’s correlation and visualized using *corrplot* (ordered by *hclust*). Heritability (h^2^) of the immunological and TB disease-related traits was calculated by estimating the percent of variation attributed to a genotype as previously described (Noll et al., 2020).

### Genotyping and QTL mapping

A subset of the inbred CC mice used in the analysis were genotyped on the GigaMUGA array (Morgan et al., 2015) available from Neogen Inc. The inbred parents, F_1_ s and F_2_ mice from the CC030xCC029 cross were genotyped on the MiniMUGA array (Sigmon et al., 2020) at Neogen Inc.

For CC030 x CC029 F_2_ analysis, markers were filtered, restricting only to those diagnostic between these strains (See **Table S3**). To assess the relative impact of the Chr7 and 15 loci, a linear regression on CFU was run with CAST haplotypes on Chr7, Chr15, and their interaction as factors.

For QTL mapping in the CC panel, the Most Recent Common Ancestor (Srivastava et al., 2017) 36-state haplotypes were downloaded from the UNC Systems Genetics Core Facility and simplified to 8-state haplotype probabilities (for the 8 CC founder strains), which is appropriate for additive genetic mapping. We generated 36-state haplotype probabilities from the individual CC mice genotyped on GigaMUGA and combined these data with the MRCA data to obtain a common genome cache.

For CC QTL analysis, genotype and phenotype data were imported into R (version 3.6.1) and reformatted for R/qtl2 (version 0.20) (Broman et al., 2019). Individual TnSeq and clinical trait phenotype values were winsorized (q = 0.006) as above. GigaMUGA annotations were downloaded from the Jackson Laboratory, and markers were thinned to a spacing of 0.1 cM using the reduce_markers function of R/qtl2. The final genetic map contained 10,067 markers. QTL mapping was carried out using a linear mixed model with LOCO (leave one chromosome out) kinship. For clinical trait scans, batch (denoted by “block” in Table S1) was included as an additive covariate. Significance thresholds for QTL were estimated using 10,000 permutations (scan1_perm function). For each trait, the maximum LOD scores from the permutation scans were used to fit generalized extreme value distributions, from which genome-wide permutation p-values were calculated. LOD profiles and effect plots were generated using the plotting functions of the R/qtl2 package. Multiple QTL at similar genetic locations were assessed for independence using qtl2pleio with 400 bootstrap samples (Boehm et al., 2019). The quantile-based permutation thresholding method of Neto et al. (Neto et al., 2012) was used to assess the statistical significance of co-mapping traits. The NL-method, which determines the LOD thresholds controlling genome wide error rate for a given p-value and “hotspot” size, was employed.

### Candidate gene prioritization

To identify potential candidate genes, we focused on three QTL that were either statistically significant (*Tip5, Hip42*) or were validated by intercross (*Tip8*). For each QTL interval (determined by Bayesian interval in qtl2), we identified mouse genes that were in differentially expressed modules between infected lungs of resistant and susceptible mouse strains (Moreira-Teixeira et al., 2020). Of these genes, we next used the Sanger sequence data (Keane et al., 2011) to filter on genetic variants segregating between CC haplotypes. Where there were clear causal haplotypes, we further filtered to genes with missense or nonsense variants.

## Data availability

All relevant data to support the findings of this study are located within the paper and supplemental files or are in the process of being made publicly available at the time of this submission; all mouse phenotype data are being deposited in Mouse Phenome Database (https://phenome.jax.org), and sequence data is being deposited in the NCBI Gene Expression Omnibus (GEO).

## Supplemental information titles and legends

**Table S1 - CC TB phenotypes**. TB disease-related phenotypes measured in the CC and parental strains. Recorded values are the average of measurements from 2-6 mice per genotype (average n=3). Mice were infected over 3 batches (denoted by “block”). “Blaze” denotes genotypes with white head-spotting coat color trait (WSB haplotype for *Kitl*; used as a positive control/proof-of-concept for QTL mapping as per (Aylor et al., 2011; Smith et al., 2019).

**Table S2 - Heritability (h**^**2**^**) estimates for each measured TB-disease associated phenotype (Tuberculosis ImmunoPhenotypes)**. h^2^ was calculated from the percentage of variation attributed to strain differences in each trait across the CC strains, as previously described (Noll et al., 2020). Weight change is the percentage of weight (grams), CFU/CEQ is log_10_ transformed, cytokines are measured in pg/mL lung homogenate and log_10_ transformed.

**Table S3 - F**_**2**_ **Intercross phenotype data**. Lung CFU measured in 46 F_2_ mice derived from CC030xCC029 intercross strategy. F_2_ mice were CAST at chromosome 7, CAST at Chromosome 15, CAST at both or CAST at neither locus. The infected F_2_ cohort included both male and female mice, as indicated.

**Table S4 - TnSeq Summary Table**. LFC values represent the log_2_ fold-change (LFC) between input and mouse-selected pools. “NA” indicates genes with fewer than 3 occupied TA transposon insertion sites for the indicated comparison. Qvals represent adjusted p-values comparing mutant abundance in input and selected pools. “NA” indicates genes with fewer than 3 occupied TA transposon insertion sites for the indicated comparison. Required *in vivo*: “TRUE” indicates the mutant is significantly underrepresented (Qval<0.05) after in mouse-selection in at least two mouse strains. Required in B6: “TRUE” indicates the mutant is significantly underrepresented (Qval <0.05) after in selection in B6 mice. Required in KO mice: “TRUE” indicates the mutant is significantly underrepresented (Qval<0.05) after in selection in *Rag*^-/^-, *Nos2*^-/-^, *Cybb*^-/-^, or *Ifng*^-/-^ mice. Core gene set: TRUE indicates the mutant is significantly underrepresented (Qval<0.05) in 30 mouse strains. “Module” corresponds to WGCNA module number as illustrated in **Figure 5A**. Mouse strains are listed in the same order as **Figure 5B**, with the corresponding cluster designation.

**Table S5 - *Hip*QTL for single *Mtb* mutant QTL and eigentrait/module QTL**. *Hip1-41* each represent host loci associated with the relative abundance of a single mutant (p<0.05). *Hip42-46* correspond to *Mtb* eigentraits identified in network analysis in **Figure 5** (including significant p<0.05 and suggestive p<0.25). Figure column headings: QTL, quantitative trait loci; *Mtb, Mycobacterium tuberculosis*; Module #, number determined from WGCNA modules; ORF, open reading frame; ID, identification number; LOD, logarithmic of the odds; Chr, chromosome.

**Table S6 - Candidate genes within QTL regions**. Prioritized candidates shown for selected QTL. Candidates were prioritized by filtering on 1) differential expression during *Mtb* infection, and 2) variants within TB-expressed genes that segregated between informative CC haplotypes. Genes listed below contain non-synonymous variants (ie. amino acid changes, regulatory mutations or splicing mutations) consistent with the identified singly causal haplotype (NOD for *Tip5;* CAST for *Tip8*). *Hip42* displayed a more complex haplotype pattern (WSB/NOD vs AJ/B6/NZO), and candidate selection is discussed in the main text. Genes with missense or nonsense variants (denoted by ∗).

**Figure S1 - Phenotypic relationships between TB disease-related traits**. Correlation between 32-measured TB traits was determined by Pearson’s correlation and visualized using corrplot version 0.84 (ordered by hclust method “complete”) in R version 4.0.3. Violet indicates a positive correlation, and yellow indicates negative correlations. The correlation coefficient for each trait comparison (r value) is noted on each square. EoD (earliness of Death); CEQ (Chromosomal equivalents).

**Figure S2 - Visual representation of QTL mapped in the CC TnSeq infection screen**. Tuberculosis ImmunoPhenotypes (*Tip*) QTL (QTL mapped by disease-associated traits in CC mice), are shown in green. *Tip*QTL mapped by separate traits that share similar founder effects were considered to be the same QTL and were named accordingly. Host Interacting with Pathogen (*Hip*) QTL, (QTL mapped by individual TnSeq mutant relative abundance profiles), are shown in purple. After WGCNA mutant clustering and mapping with representative eigengenes from each module, QTL mapped by module eigengenes are shown in magenta.

**Figure S3 - Module-trait associations**. Rows correspond to modules, columns to clinical traits. Numbers in each cell give the Pearson correlation between the module eigengene and the trait values across the 60-mouse panel (p-values in parentheses). Cells are colored by correlation as shown in the color legend (right).

## References

Abel, L., Fellay, J., Haas, D.W., Schurr, E., Srikrishna, G., Urbanowski, M., Chaturvedi, N., Srinivasan, S., Johnson, D.H., and Bishai, W.R. (2018). Genetics of human susceptibility to active and latent tuberculosis: present knowledge and future perspectives. Lancet Infect Dis 18, e64–e75.

Adams, L.A., Möller, M., Nebel, A., Schreiber, S., van der Merwe, L., van Helden, P.D., and Hoal, E.G. (2011). Polymorphisms in MC3R promoter and CTSZ 3’UTR are associated with tuberculosis susceptibility. Eur J Hum Genet 19, 676–681.

Ahmed, M., Thirunavukkarasu, S., Rosa, B.A., Thomas, K.A., Das, S., Rangel-Moreno, J., Lu, L., Mehra, S., Mbandi, S.K., Thackray, L.B., et al. (2020). Immune correlates of tuberculosis disease and risk translate across species. Sci Transl Med 12.

Altare, F., Durandy, A., Lammas, D., Emile, J.F., Lamhamedi, S., Le Deist, F., Drysdale, P., Jouanguy, E., Döffinger, R., Bernaudin, F., et al. (1998). Impairment of mycobacterial immunity in human interleukin-12 receptor deficiency. Science 280, 1432–1435.

Amaral, E.P., Riteau, N., Moayeri, M., Maier, N., Mayer-Barber, K.D., Pereira, R.M., Lage, S.L., Kubler, A., Bishai, W.R., D’Império-Lima, M.R., et al. (2018). Lysosomal Cathepsin Release Is Required for NLRP3-Inflammasome Activation by Mycobacterium tuberculosis in Infected Macrophages. Front Immunol 9, 1427.

Ansari, M.A., Pedergnana, V., C, L.C.I., Magri, A., Von Delft, A., Bonsall, D., Chaturvedi, N., Bartha, I., Smith, D., Nicholson, G., et al. (2017). Genome-to-genome analysis highlights the effect of the human innate and adaptive immune systems on the hepatitis C virus. Nat Genet 49, 666–673.

Aylor, D.L., Valdar, W., Foulds-Mathes, W., Buus, R.J., Verdugo, R.A., Baric, R.S., Ferris, M.T., Frelinger, J.A., Heise, M., Frieman, M.B., et al. (2011). Genetic analysis of complex traits in the emerging Collaborative Cross. Genome Res 21, 1213–1222.

Ayres, J.S., and Schneider, D.S. (2012). Tolerance of infections. Annu Rev Immunol 30, 271–294.

Barber, D.L., Mayer-Barber, K.D., Feng, C.G., Sharpe, A.H., and Sher, A. (2011). CD4 T cells promote rather than control tuberculosis in the absence of PD-1-mediated inhibition. J Immunol 186, 1598–1607.

Bellerose, M.M., Baek, S.-H., Huang, C.-C., Moss, C.E., Koh, E.-I., Proulx, M.K., Smith, C.M., Baker, R.E., Lee, J.S., Eum, S., et al. (2019). Common Variants in the Glycerol Kinase Gene Reduce Tuberculosis Drug Efficacy. mBio 10, e00663–00619.

Bellerose, M.M., Proulx, M.K., Smith, C.M., Baker, R.E., Ioerger, T.R., and Sassetti, C.M. (2020). Distinct Bacterial Pathways Influence the Efficacy of Antibiotics against Mycobacterium tuberculosis. mSystems 5.

Berthenet, E., Yahara, K., Thorell, K., Pascoe, B., Meric, G., Mikhail, J.M., Engstrand, L., Enroth, H., Burette, A., Megraud, F., et al. (2018). A GWAS on Helicobacter pylori strains points to genetic variants associated with gastric cancer risk. BMC Biol 16, 84.

Boehm, F.J., Chesler, E.J., Yandell, B.S., and Broman, K.W. (2019). Testing Pleiotropy vs. Separate QTL in Multiparental Populations. G3 (Bethesda) 9, 2317–2324.

Bogunovic, D., Byun, M., Durfee, L.A., Abhyankar, A., Sanal, O., Mansouri, D., Salem, S., Radovanovic, I., Grant, A.V., Adimi, P., et al. (2012). Mycobacterial disease and impaired IFN-γ immunity in humans with inherited ISG15 deficiency. Science 337, 1684–1688.

Broman, K.W., Gatti, D.M., Simecek, P., Furlotte, N.A., Prins, P., Sen, S., Yandell, B.S., and Churchill, G.A. (2019). R/qtl2: Software for Mapping Quantitative Trait Loci with High-Dimensional Data and Multiparent Populations. Genetics 211, 495–502.

Bustamante, J., Boisson-Dupuis, S., Abel, L., and Casanova, J.L. (2014). Mendelian susceptibility to mycobacterial disease: genetic, immunological, and clinical features of inborn errors of IFN-γ immunity. Semin Immunol 26, 454–470.

Caruso, A.M., Serbina, N., Klein, E., Triebold, K., Bloom, B.R., and Flynn, J.L. (1999). Mice deficient in CD4 T cells have only transiently diminished levels of IFN-gamma, yet succumb to tuberculosis. J Immunol 162, 5407–5416.

Caws, M., Thwaites, G., Dunstan, S., Hawn, T.R., Lan, N.T., Thuong, N.T., Stepniewska, K., Huyen, M.N., Bang, N.D., Loc, T.H., et al. (2008). The influence of host and bacterial genotype on the development of disseminated disease with Mycobacterium tuberculosis. PLoS Pathog 4, e1000034.

Chen, L., Xie, Q.W., and Nathan, C. (1998). Alkyl hydroperoxide reductase subunit C (AhpC) protects bacterial and human cells against reactive nitrogen intermediates. Mol Cell 1, 795–805.

Churchill, G.A., Airey, D.C., Allayee, H., Angel, J.M., Attie, A.D., Beatty, J., Beavis, W.D., Belknap, J.K., Bennett, B., Berrettini, W., et al. (2004). The Collaborative Cross, a community resource for the genetic analysis of complex traits. Nat Genet 36, 1133–1137.

Churchill, G.A., Gatti, D.M., Munger, S.C., and Svenson, K.L. (2012). The Diversity Outbred mouse population. Mamm Genome 23, 713–718.

Comstock, G.W. (1978). Tuberculosis in twins: a re-analysis of the Prophit survey. Am Rev Respir Dis 117, 621–624.

Cooke, G.S., Campbell, S.J., Bennett, S., Lienhardt, C., McAdam, K.P., Sirugo, G., Sow, O., Gustafson, P., Mwangulu, F., van Helden, P., et al. (2008). Mapping of a novel susceptibility locus suggests a role for MC3R and CTSZ in human tuberculosis. Am J Respir Crit Care Med 178, 203–207.

Cooper, A.M., Dalton, D.K., Stewart, T.A., Griffin, J.P., Russell, D.G., and Orme, I.M. (1993). Disseminated tuberculosis in interferon gamma gene-disrupted mice. J Exp Med 178, 2243–2247.

Cooper, A.M., Magram, J., Ferrante, J., and Orme, I.M. (1997). Interleukin 12 (IL-12) is crucial to the development of protective immunity in mice intravenously infected with mycobacterium tuberculosis. J Exp Med 186, 39–45.

de Keijzer, J., de Haas, P.E., de Ru, A.H., van Veelen, P.A., and van Soolingen, D. (2014). Disclosure of selective advantages in the “modern” sublineage of the Mycobacterium tuberculosis Beijing genotype family by quantitative proteomics. Mol Cell Proteomics 13, 2632–2645.

DeJesus, M.A., Ambadipudi, C., Baker, R., Sassetti, C., and Ioerger, T.R. (2015). TRANSIT--A Software Tool for Himar1 TnSeq Analysis. PLoS Comput Biol 11, e1004401.

Dick, T., Manjunatha, U., Kappes, B., and Gengenbacher, M. (2010). Vitamin B6 biosynthesis is essential for survival and virulence of Mycobacterium tuberculosis. Mol Microbiol 78, 980–988.

Ferris, M.T., Aylor, D.L., Bottomly, D., Whitmore, A.C., Aicher, L.D., Bell, T.A., Bradel-Tretheway, B., Bryan, J.T., Buus, R.J., Gralinski, L.E., et al. (2013). Modeling host genetic regulation of influenza pathogenesis in the collaborative cross. PLoS Pathog 9, e1003196.

Filipe-Santos, O., Bustamante, J., Haverkamp, M.H., Vinolo, E., Ku, C.L., Puel, A., Frucht, D.M., Christel, K., von Bernuth, H., Jouanguy, E., et al. (2006). X-linked susceptibility to mycobacteria is caused by mutations in NEMO impairing CD40-dependent IL-12 production. J Exp Med 203, 1745–1759.

Flynn, J.L., Chan, J., Triebold, K.J., Dalton, D.K., Stewart, T.A., and Bloom, B.R. (1993). An essential role for interferon gamma in resistance to Mycobacterium tuberculosis infection. J Exp Med 178, 2249–2254.

Gagneux, S., DeRiemer, K., Van, T., Kato-Maeda, M., de Jong, B.C., Narayanan, S., Nicol, M., Niemann, S., Kremer, K., Gutierrez, M.C., et al. (2006). Variable host-pathogen compatibility in Mycobacterium tuberculosis. Proc Natl Acad Sci U S A 103, 2869–2873.

Geiman, D.E., Kaushal, D., Ko, C., Tyagi, S., Manabe, Y.C., Schroeder, B.G., Fleischmann, R.D., Morrison, N.E., Converse, P.J., Chen, P., et al. (2004). Attenuation of late-stage disease in mice infected by the Mycobacterium tuberculosis mutant lacking the SigF alternate sigma factor and identification of SigF-dependent genes by microarray analysis. Infect Immun 72, 1733–1745.

Goitre, L., Trapani, E., Trabalzini, L., and Retta, S.F. (2014). The Ras superfamily of small GTPases: the unlocked secrets. Methods Mol Biol 1120, 1–18.

Gonzalo-Asensio, J., Malaga, W., Pawlik, A., Astarie-Dequeker, C., Passemar, C., Moreau, F., Laval, F., Daffé, M., Martin, C., Brosch, R., et al. (2014). Evolutionary history of tuberculosis shaped by conserved mutations in the PhoPR virulence regulator. Proc Natl Acad Sci U S A 111, 11491–11496.

Gopal, R., Lin, Y., Obermajer, N., Slight, S., Nuthalapati, N., Ahmed, M., Kalinski, P., and Khader, S.A. (2012). IL-23-dependent IL-17 drives Th1-cell responses following Mycobacterium bovis BCG vaccination. European journal of immunology 42, 364 – 373.

Gopal, R., Monin, L., Torres, D., Slight, S., Mehra, S., McKenna, K.C., Fallert Junecko, B.A., Reinhart, T.A., Kolls, J., Báez-Saldaña, R., et al. (2013). S100A8/A9 proteins mediate neutrophilic inflammation and lung pathology during tuberculosis. Am J Respir Crit Care Med 188, 1137–1146.

Gralinski, L.E., Menachery, V.D., Morgan, A.P., Totura, A.L., Beall, A., Kocher, J., Plante, J., Harrison-Shostak, D.C., Schäfer, A., Pardo-Manuel de Villena, F., et al. (2017). Allelic Variation in the Toll-Like Receptor Adaptor Protein Ticam2 Contributes to SARS-Coronavirus Pathogenesis in Mice. G3 (Bethesda) 7, 1653–1663.

Hillas, P.J., del Alba, F.S., Oyarzabal, J., Wilks, A., and Ortiz De Montellano, P.R. (2000). The AhpC and AhpD antioxidant defense system of Mycobacterium tuberculosis. J Biol Chem 275, 18801–18809.

Holt, K.E., McAdam, P., Thai, P.V.K., Thuong, N.T.T., Ha, D.T.M., Lan, N.N., Lan, N.H., Nhu, N.T.Q., Hai, H.T., Ha, V.T.N., et al. (2018). Frequent transmission of the Mycobacterium tuberculosis Beijing lineage and positive selection for the EsxW Beijing variant in Vietnam. Nat Genet 50, 849–856.

Keane, T.M., Goodstadt, L., Danecek, P., White, M.A., Wong, K., Yalcin, B., Heger, A., Agam, A., Slater, G., Goodson, M., et al. (2011). Mouse genomic variation and its effect on phenotypes and gene regulation. Nature 477, 289–294.

Keller, M.P., Gatti, D.M., Schueler, K.L., Rabaglia, M.E., Stapleton, D.S., Simecek, P., Vincent, M., Allen, S., Broman, A.T., Bacher, R., et al. (2018). Genetic Drivers of Pancreatic Islet Function. Genetics 209, 335–356.

Khader, S.A., Bell, G.K., Pearl, J.E., Fountain, J.J., Rangel-Moreno, J., Cilley, G.E., Shen, F., Eaton, S.M., Gaffen, S.L., Swain, S.L., et al. (2007). IL-23 and IL-17 in the establishment of protective pulmonary CD4+ T cell responses after vaccination and during Mycobacterium tuberculosis challenge. Nat Immunol 8, 369–377.

Langfelder, P., and Horvath, S. (2008). WGCNA: an R package for weighted correlation network analysis. Bmc Bioinformatics 9, 559.

Lázár-Molnár, E., Chen, B., Sweeney, K.A., Wang, E.J., Liu, W., Lin, J., Porcelli, S.A., Almo, S.C., Nathenson, S.G., and Jacobs, W.R., Jr. (2010). Programmed death-1 (PD-1)-deficient mice are extraordinarily sensitive to tuberculosis. Proc Natl Acad Sci U S A 107, 13402–13407.

Le Chevalier, F., Cascioferro, A., Frigui, W., Pawlik, A., Boritsch, E.C., Bottai, D., Majlessi, L., Herrmann, J.L., and Brosch, R. (2015). Revisiting the role of phospholipases C in virulence and the lifecycle of Mycobacterium tuberculosis. Sci Rep 5, 16918.

Lin, P.L., Ford, C.B., Coleman, M.T., Myers, A.J., Gawande, R., Ioerger, T., Sacchettini, J., Fortune, S.M., and Flynn, J.L. (2014). Sterilization of granulomas is common in active and latent tuberculosis despite within-host variability in bacterial killing. Nat Med 20, 75–79.

Long, J.E., DeJesus, M., Ward, D., Baker, R.E., Ioerger, T., and Sassetti, C.M. (2015). Identifying essential genes in Mycobacterium tuberculosis by global phenotypic profiling. Methods Mol Biol 1279, 79–95.

Long, M., Kranjc, T., Mysior, M.M., and Simpson, J.C. (2020). RNA Interference Screening Identifies Novel Roles for RhoBTB1 and RhoBTB3 in Membrane Trafficking Events in Mammalian Cells. Cells 9.

Lu, L.L., Smith, M.T., Yu, K.K.Q., Luedemann, C., Suscovich, T.J., Grace, P.S., Cain, A., Yu, W.H., McKitrick, T.R., Lauffenburger, D., et al. (2019). IFN-γ-independent immune markers of Mycobacterium tuberculosis exposure. Nat Med 25, 977–987.

Martin, C.J., Cadena, A.M., Leung, V.W., Lin, P.L., Maiello, P., Hicks, N., Chase, M.R., Flynn, J.L., and Fortune, S.M. (2017). Digitally Barcoding Mycobacterium tuberculosis Reveals In Vivo Infection Dynamics in the Macaque Model of Tuberculosis. mBio 8.

McHenry, M.L., Williams, S.M., and Stein, C.M. (2020). Genetics and evolution of tuberculosis pathogenesis: New perspectives and approaches. Infect Genet Evol 81, 104204.

Minch, K.J., Rustad, T.R., Peterson, E.J., Winkler, J., Reiss, D.J., Ma, S., Hickey, M., Brabant, W., Morrison, B., Turkarslan, S., et al. (2015). The DNA-binding network of Mycobacterium tuberculosis. Nat Commun 6, 5829.

Mishra, B.B., Lovewell, R.R., Olive, A.J., Zhang, G.L., Wang, W.F., Eugenin, E., Smith, C.M., Phuah, J.Y., Long, J.E., Dubuke, M.L., et al. (2017). Nitric oxide prevents a pathogen-permissive granulocytic inflammation during tuberculosis. Nature Microbiology 2.

Mishra, B.B., Rathinam, V.A., Martens, G.W., Martinot, A.J., Kornfeld, H., Fitzgerald, K.A., and Sassetti, C.M. (2013). Nitric oxide controls the immunopathology of tuberculosis by inhibiting NLRP3 inflammasome-dependent processing of IL-1β. Nat Immunol 14, 52–60.

Moreira-Teixeira, L., Tabone, O., Graham, C.M., Singhania, A., Stavropoulos, E., Redford, P.S., Chakravarty, P., Priestnall, S.L., Suarez-Bonnet, A., Herbert, E., et al. (2020). Mouse transcriptome reveals potential signatures of protection and pathogenesis in human tuberculosis. Nat Immunol 21, 464–476.

Morgan, A.P., Fu, C.P., Kao, C.Y., Welsh, C.E., Didion, J.P., Yadgary, L., Hyacinth, L., Ferris, M.T., Bell, T.A., Miller, D.R., et al. (2015). The Mouse Universal Genotyping Array: From Substrains to Subspecies. G3 (Bethesda) 6, 263–279.

Morth, J.P., Gosmann, S., Nowak, E., and Tucker, P.A. (2005). A novel two-component system found in Mycobacterium tuberculosis. FEBS Lett 579, 4145–4148.

Muñoz-Elías, E.J., Timm, J., Botha, T., Chan, W.T., Gomez, J.E., and McKinney, J.D. (2005). Replication dynamics of Mycobacterium tuberculosis in chronically infected mice. Infect Immun 73, 546–551.

Murphy, K.C., Nelson, S.J., Nambi, S., Papavinasasundaram, K., Baer, C.E., and Sassetti, C.M. (2018). ORBIT: a New Paradigm for Genetic Engineering of Mycobacterial Chromosomes. mBio 9.

Nambi, S., Long, J.E., Mishra, B.B., Baker, R., Murphy, K.C., Olive, A.J., Nguyen, H.P., Shaffer, S.A., and Sassetti, C.M. (2015). The Oxidative Stress Network of Mycobacterium tuberculosis Reveals Coordination between Radical Detoxification Systems. Cell Host Microbe 17, 829–837.

Neto, E.C., Keller, M.P., Broman, A.F., Attie, A.D., Jansen, R.C., Broman, K.W., and Yandell, B.S. (2012). Quantile-based permutation thresholds for quantitative trait loci hotspots. Genetics 191, 1355–1365.

Niazi, M.K., Dhulekar, N., Schmidt, D., Major, S., Cooper, R., Abeijon, C., Gatti, D.M., Kramnik, I., Yener, B., Gurcan, M., et al. (2015). Lung necrosis and neutrophils reflect common pathways of susceptibility to Mycobacterium tuberculosis in genetically diverse, immune-competent mice. Dis Model Mech 8, 1141–1153.

Noll, K.E., Ferris, M.T., and Heise, M.T. (2019). The Collaborative Cross: A Systems Genetics Resource for Studying Host-Pathogen Interactions. Cell Host Microbe 25, 484–498.

Noll, K.E., Whitmore, A.C., West, A., McCarthy, M.K., Morrison, C.R., Plante, K.S., Hampton, B.K., Kollmus, H., Pilzner, C., Leist, S.R., et al. (2020). Complex Genetic Architecture Underlies Regulation of Influenza-A-Virus-Specific Antibody Responses in the Collaborative Cross. Cell Rep 31, 107587.

Obermajer, N., Svajger, U., Bogyo, M., Jeras, M., and Kos, J. (2008). Maturation of dendritic cells depends on proteolytic cleavage by cathepsin X. J Leukoc Biol 84, 1306–1315.

Olive, A.J., Smith, C.M., Kiritsy, M.C., and Sassetti, C.M. (2018). The Phagocyte Oxidase Controls Tolerance to Mycobacterium tuberculosis Infection. J Immunol 201, 1705–1716.

Palace, S.G., Proulx, M.K., Lu, S., Baker, R.E., and Goguen, J.D. (2014). Genome-wide mutant fitness profiling identifies nutritional requirements for optimal growth of Yersinia pestis in deep tissue. mBio 5.

Pandey, A.K., and Sassetti, C.M. (2008). Mycobacterial persistence requires the utilization of host cholesterol. Proc Natl Acad Sci U S A 105, 4376–4380.

Pethe, K., Sequeira, P.C., Agarwalla, S., Rhee, K., Kuhen, K., Phong, W.Y., Patel, V., Beer, D., Walker, J.R., Duraiswamy, J., et al. (2010). A chemical genetic screen in Mycobacterium tuberculosis identifies carbon-source-dependent growth inhibitors devoid of in vivo efficacy. Nat Commun 1, 57.

Rodrigue, S., Provvedi, R., Jacques, P.E., Gaudreau, L., and Manganelli, R. (2006). The sigma factors of Mycobacterium tuberculosis. FEMS Microbiol Rev 30, 926–941.

Ryndak, M.B., Wang, S., Smith, I., and Rodriguez, G.M. (2010). The Mycobacterium tuberculosis high-affinity iron importer, IrtA, contains an FAD-binding domain. J Bacteriol 192, 861–869.

Sakai, S., Kauffman, K.D., Sallin, M.A., Sharpe, A.H., Young, H.A., Ganusov, V.V., and Barber, D.L. (2016). CD4 T Cell-Derived IFN-γ Plays a Minimal Role in Control of Pulmonary Mycobacterium tuberculosis Infection and Must Be Actively Repressed by PD-1 to Prevent Lethal Disease. PLoS Pathog 12, e1005667.

Sassetti, C.M., Boyd, D.H., and Rubin, E.J. (2003). Genes required for mycobacterial growth defined by high density mutagenesis. Mol Microbiol 48, 77–84.

Sassetti, C.M., and Rubin, E.J. (2003). Genetic requirements for mycobacterial survival during infection. Proc Natl Acad Sci U S A 100, 12989–12994.

Saul, M.C., Philip, V.M., Reinholdt, L.G., and Chesler, E.J. (2019). High-Diversity Mouse Populations for Complex Traits. Trends Genet 35, 501–514.

Saunders, B.M., Frank, A.A., Orme, I.M., and Cooper, A.M. (2002). CD4 is required for the development of a protective granulomatous response to pulmonary tuberculosis. Cell Immunol 216, 65–72.

Shorter, J.R., Najarian, M.L., Bell, T.A., Blanchard, M., Ferris, M.T., Hock, P., Kashfeen, A., Kirchoff, K.E., Linnertz, C.L., Sigmon, J.S., et al. (2019). Whole Genome Sequencing and Progress Toward Full Inbreeding of the Mouse Collaborative Cross Population. G3 (Bethesda) 9, 1303–1311.

Sigmon, J.S., Blanchard, M.W., Baric, R.S., Bell, T.A., Brennan, J., Brockmann, G.A., Burks, A.W., Calabrese, J.M., Caron, K.M., Cheney, R.E., et al. (2020). Content and performance of the MiniMUGA genotyping array, a new tool to improve rigor and reproducibility in mouse research. bioRxiv, 2020.2003.2012.989400.

Smith, C.M., Proulx, M.K., Lai, R., Kiritsy, M.C., Bell, T.A., Hock, P., Pardo-Manuel de Villena, F., Ferris, M.T., Baker, R.E., Behar, S.M., et al. (2019). Functionally Overlapping Variants Control Tuberculosis Susceptibility in Collaborative Cross Mice. mBio 10.

Smith, C.M., Proulx, M.K., Olive, A.J., Laddy, D., Mishra, B.B., Moss, C., Gutierrez, N.M., Bellerose, M.M., Barreira-Silva, P., Phuah, J.Y., et al. (2016). Tuberculosis Susceptibility and Vaccine Protection Are Independently Controlled by Host Genotype. mBio 7.

Smith, C.M., and Sassetti, C.M. (2018). Modeling Diversity: Do Homogeneous Laboratory Strains Limit Discovery? Trends Microbiol 26, 892–895.

Sousa, J., Cá, B., Maceiras, A.R., Simões-Costa, L., Fonseca, K.L., Fernandes, A.I., Ramos, A., Carvalho, T., Barros, L., Magalhães, C., et al. (2020). Mycobacterium tuberculosis associated with severe tuberculosis evades cytosolic surveillance systems and modulates IL-1β production. Nat Commun 11, 1949.

Springer, B., Master, S., Sander, P., Zahrt, T., McFalone, M., Song, J., Papavinasasundaram, K.G., Colston, M.J., Boettger, E., and Deretic, V. (2001). Silencing of oxidative stress response in Mycobacterium tuberculosis: expression patterns of ahpC in virulent and avirulent strains and effect of ahpC inactivation. Infect Immun 69, 5967–5973.

Srivastava, A., Morgan, A.P., Najarian, M.L., Sarsani, V.K., Sigmon, J.S., Shorter, J.R., Kashfeen, A., McMullan, R.C., Williams, L.H., Giusti-Rodríguez, P., et al. (2017). Genomes of the Mouse Collaborative Cross. Genetics 206, 537–556.

Stanley, S.A., Raghavan, S., Hwang, W.W., and Cox, J.S. (2003). Acute infection and macrophage subversion by Mycobacterium tuberculosis require a specialized secretion system. Proceedings of the National Academy of Sciences 100, 13001–13006.

Svenson, K.L., Gatti, D.M., Valdar, W., Welsh, C.E., Cheng, R., Chesler, E.J., Palmer, A.A., McMillan, L., and Churchill, G.A. (2012). High-resolution genetic mapping using the Mouse Diversity outbred population. Genetics 190, 437–447.

Thuong, N.T., Tram, T.T., Dinh, T.D., Thai, P.V., Heemskerk, D., Bang, N.D., Chau, T.T., Russell, D.G., Thwaites, G.E., Hawn, T.R., et al. (2016). MARCO variants are associated with phagocytosis, pulmonary tuberculosis susceptibility and Beijing lineage. Genes Immun 17, 419–425.

Turkarslan, S., Peterson, E.J., Rustad, T.R., Minch, K.J., Reiss, D.J., Morrison, R., Ma, S., Price, N.D., Sherman, D.R., and Baliga, N.S. (2015). A comprehensive map of genome-wide gene regulation in Mycobacterium tuberculosis. Sci Data 2, 150010.

Wang, Q., Boshoff, H.I.M., Harrison, J.R., Ray, P.C., Green, S.R., Wyatt, P.G., and Barry, C.E., 3rd (2020). PE/PPE proteins mediate nutrient transport across the outer membrane of Mycobacterium tuberculosis. Science 367, 1147–1151.

Woong Park, S., Klotzsche, M., Wilson, D.J., Boshoff, H.I., Eoh, H., Manjunatha, U., Blumenthal, A., Rhee, K., Barry, C.E., 3rd, Aldrich, C.C., et al. (2011). Evaluating the sensitivity of Mycobacterium tuberculosis to biotin deprivation using regulated gene expression. PLoS Pathog 7, e1002264.

Zak, D.E., Penn-Nicholson, A., Scriba, T.J., Thompson, E., Suliman, S., Amon, L.M., Mahomed, H., Erasmus, M., Whatney, W., Hussey, G.D., et al. (2016). A blood RNA signature for tuberculosis disease risk: a prospective cohort study. Lancet 387, 2312–2322.

Zhang, J., Teh, M., Kim, J., Eva, M.M., Cayrol, R., Meade, R., Nijnik, A., Montagutelli, X., Malo, D., and Jaubert, J. (2019). A Loss-of-Function Mutation in the Integrin Alpha L (Itgal) Gene Contributes to Susceptibility to Salmonella enterica Serovar Typhimurium Infection in Collaborative Cross Strain CC042. Infect Immun 88.

Zhang, Y.J., Reddy, M.C., Ioerger, T.R., Rothchild, A.C., Dartois, V., Schuster, B.M., Trauner, A., Wallis, D., Galaviz, S., Huttenhower, C., et al. (2013). Tryptophan Biosynthesis Protects Mycobacteria from CD4 T-Cell-Mediated Killing. Cell 155, 1296 – 1308.

