## Supplementary material for "Host-pathogen genetic interactions underlie tuberculosis susceptibility": TableS2_h2_TIPs.pdf

**Table S2. Heritability ($h^2$) estimates for each measured clinical trait (Tuberculosis ImmunoPhenotypes; Tip)**. $h^2$ was calculated from the percentage of variation attributed to strain differences in each trait across the CC strains, as previously described (Noll et al. 2020). Weight change is the percentage of weight (grams), CFU/CEQ is $log_{10}$ transformed, cytokines are measured in pg/mL lung homogenate and $log_{10}$ transformed**.**

| **Trait** | **$h^2$ (%)** |
|---|---|
| Weight change | 67.50 |
| Spleen CFU | 72.90 |
| Spleen CEQ | 60.40 |
| Lung CFU | 83.30 |
| Lung CEQ | 80.60 |
| Eotaxin | 64.20 |
| G-CSF | 77.80 |
| IFNy | 87.70 |
| IL-1a | 75.60 |
| IL-1B | 71.70 |
| IL-2 | 62.30 |
| IL-6 | 73.90 |
| IL-9 | 56.80 |
| IL-10 | 64.00 |
| IL-17 | 78.80 |
| IP-10 | 80.10 |
| CXCL1 | 82.00 |
| LIF | 73.40 |
| LIX | 65.20 |
| MCP-1 | 77.00 |
| M-CSF | 79.30 |
| MIG | 81.10 |
| MIP-1a | 78.00 |
| MIP-1b | 68.50 |
| MIP-2 | 80.80 |
| RANTES | 73.70 |
| TNFa | 66.30 |
