## Supplementary material for "Host-pathogen genetic interactions underlie tuberculosis susceptibility": TableS5_hipQTL.pdf

**Table S5. *Hip*QTL for single *Mtb* mutant QTL and eigentrait/module QTL**. Hip1-41 each represent host loci associated with the fitness profile of a single mutant (p<0.05). Hip42-46 correspond to *Mtb* eigentraits identified in network analysis in **Figure 5** (including significant p<0.05 and suggestive p<0.25). Figure Column headings: QTL, quantitative trait loci; *Mtb*, *Mycobacterium tuberculosis*; Module #, number determined from WGCNA modules, ORF, open reading frame; ID, identification number; LOD, logarithmic of the odds; Chr, chromosome.

| QTL | Trait | *Mtb* ORF ID | Module # | LOD | P value | Chr | Start (Mb) | Peak (Mb) | End (Mb) |
|---|---|---|---|---|---|---|---|---|---|
| *Hip1* | rv0770 | RVBD_0770 | mod17 | 9.81 | 5.61E-03 | 1 | 40.43 | 42.73 | 43.32 |
| *Hip2* | rv0309 | RVBD_0309 | mod13 | 7.95 | 3.22E-02 | 1 | 57.99 | 58.18 | 62.79 |
| *Hip3* | rv3657c | RVBD_3657c | mod15 | 7.90 | 4.95E-02 | 1 | 136.39 | 138.24 | 143.60 |
| *Hip4* | rv0110 | RVBD_0110 | mod18 | 7.79 | 4.39E-02 | 2 | 170.67 | 174.00 | 178.84 |
| *Hip5* | rv3577 | RVBD_3577 | mod7 | 9.23 | 3.83E-02 | 3 | 3.32 | 10.03 | 14.67 |
| *Hip6* | rv3005c | RVBD_3005c | mod6 | 8.03 | 3.75E-02 | 3 | 20.31 | 26.12 | 26.12 |
| *Hip7* | dinX | RVBD_1537 | mod15 | 9.97 | 5.38E-03 | 3 | 26.99 | 30.29 | 33.85 |
| *Hip8* | fadA6 | RVBD_3556c | mod5 | 8.74 | 1.01E-02 | 3 | 29.23 | 35.22 | 37.11 |
| *Hip9* | dinX | RVBD_1537 | mod15 | 9.22 | 1.60E-02 | 3 | 36.22 | 36.83 | 38.27 |
| *Hip10* | rv2707 | RVBD_2707 | mod6 | 8.17 | 4.21E-02 | 3 | 100.90 | 103.23 | 115.82 |
| *Hip11* | rv3701c | RVBD_3701c | mod6 | 7.90 | 3.38E-02 | 4 | 74.00 | 78.25 | 87.00 |
| *Hip12* | ahpC | RVBD_2428 | mod13 | 8.12 | 2.14E-02 | 6 | 19.75 | 22.21 | 23.31 |
| *Hip13* | umaA | RVBD_0469 | mod20 | 8.32 | 2.31E-02 | 7 | 117.87 | 118.41 | 120.15 |
| *Hip14* | rv2566 | RVBD_2566 | mod15 | 7.86 | 4.55E-02 | 7 | 123.21 | 126.67 | 126.67 |
| *Hip15* | rv3173c | RVBD_3173c | mod5 | 8.17 | 3.03E-02 | 7 | 137.41 | 138.36 | 138.36 |
| *Hip16* | rv3173c | RVBD_3173c | mod5 | 8.12 | 3.28E-02 | 7 | 139.15 | 140.76 | 141.88 |
| *Hip17* | rv3502c | RVBD_3502c | mod5 | 8.16 | 3.17E-02 | 9 | 15.91 | 16.33 | 18.72 |
| *Hip18* | mycP1 | RVBD_3883c | mod3 | 9.09 | 3.66E-03 | 9 | 28.47 | 29.45 | 31.10 |
| *Hip19* | rv0057 | RVBD_0057 | mod6 | 8.39 | 3.79E-02 | 9 | 36.78 | 40.07 | 40.36 |
| *Hip20* | hycE | RVBD_0087 | mod20 | 8.21 | 1.40E-02 | 9 | 47.40 | 47.93 | 51.80 |
| *Hip21* | mbtA | RVBD_2384 | mod4 | 8.30 | 2.05E-02 | 10 | 64.48 | 68.09 | 75.42 |
| *Hip22* | eccD1 | RVBD_3877 | mod3 | 8.08 | 3.08E-02 | 10 | 64.56 | 68.12 | 71.04 |
| *Hip23* | rv2989 | RVBD_2989 | mod12 | 9.16 | 1.67E-02 | 10 | 74.30 | 77.63 | 81.03 |
| *Hip24* | mce4A | RVBD_3499c | mod16 | 7.91 | 4.12E-02 | 10 | 78.88 | 81.36 | 88.25 |
| *Hip25* | treS | RVBD_0126 | mod7 | 7.94 | 3.04E-02 | 11 | 20.80 | 36.14 | 44.06 |
| *Hip26* | pckA | RVBD_0211 | mod3 | 7.67 | 4.74E-02 | 11 | 85.95 | 89.78 | 91.75 |
| *Hip27* | aspB | RVBD_3565 | mod7 | 8.32 | 3.66E-02 | 11 | 114.69 | 116.99 | 117.08 |
| *Hip28* | rv1227c | RVBD_1227c | mod17 | 9.16 | 4.64E-02 | 12 | 25.23 | 25.23 | 28.54 |
| *Hip29* | rv0219 | RVBD_0219 | mod20 | 7.94 | 3.09E-02 | 12 | 40.65 | 42.65 | 47.22 |
| *Hip30* | rv3643 | RVBD_3643 | mod8 | 8.89 | 1.04E-02 | 13 | 95.43 | 97.08 | 97.79 |
| *Hip31* | ansA | RVBD_1538c | mod11 | 8.28 | 3.32E-02 | 13 | 96.82 | 97.79 | 99.09 |
| *Hip32* | echA19 | RVBD_3516 | mod20 | 9.68 | 3.95E-02 | 13 | 113.20 | 114.59 | 117.64 |
| *Hip33* | rv1836c | RVBD_1836c | mod15 | 9.19 | 1.75E-02 | 14 | 74.94 | 76.40 | 76.43 |

| | | | | | | | | | |
|---|---|---|---|---|---|---|---|---|---|
| *Hip34* | rv2183c | RVBD_2183c | mod11 | 7.75 | 4.84E-02 | 16 | 12.18 | 14.06 | 17.92 |
| *Hip35* | rv1178 | RVBD_1178 | mod6 | 8.19 | 4.76E-02 | 17 | 80.92 | 80.92 | 83.23 |
| *Hip36* | rv0492c | RVBD_0492c | mod17 | 8.90 | 3.18E-02 | 18 | 5.85 | 5.85 | 12.40 |
| *Hip37* | cysM | RVBD_1336 | mod12 | 8.47 | 8.67E-03 | 19 | 4.20 | 6.46 | 6.46 |
| *Hip38* | atsA | RVBD_0711 | mod1 | 8.58 | 1.38E-02 | 19 | 31.21 | 37.86 | 37.93 |
| *Hip39* | galE2 | RVBD_0501 | mod6 | 8.10 | 2.85E-02 | X | 6.01 | 6.01 | 9.12 |
| *Hip40* | pks11 | RVBD_1665 | mod17 | 8.25 | 1.94E-02 | X | 50.43 | 51.75 | 52.29 |
| *Hip41* | pknK | RVBD_3080c | mod17 | 8.73 | 3.79E-02 | X | 95.01 | 102.02 | 130.04 |
| *Hip42* | Module 3 | ESX1 operon | mod3 | 7.80 | 5.38E-02 | 10 | 64.7 | 68.27 | 77.07 |
| *Hip42* | Module 4 | Mycobactin (*mbt*) | mod4 | 7.79 | 5.05E-02 | 10 | 65.23 | 69.94 | 74.30 |
| *Hip43* | Module 16 | *mce4* operon | mod16 | 7.53 | 7.97E-02 | 10 | 74.30 | 81.36 | 87.61 |
| *Hip44* | Module 19 | unclassified | Module 19 | 7.64 | 1.39E-01 | 11 | 60.87 | 62.20 | 63.26 |
| *Hip45* | Module 10 | Transcriptional regulation | Module 10 | 6.95 | 1.04E-01 | 15 | 100.39 | 102.25 | 103.36 |
| *Hip46* | Module 10 | Transcriptional regulation | Module 10 | 6.32 | 2.54E-01 | 19 | 32.74 | 32.87 | 37.48 |
