## Supplementary material for "Host-pathogen genetic interactions underlie tuberculosis susceptibility": TableS6_CandidateGenes.pdf

**Table S6. Candidate genes within QTL regions**. Prioritized candidates shown for selected QTL. Candidates were prioritized by filtering on 1) differential expression during *Mtb* infection, and 2) variants within TB-expressed genes that segregated between informative CC haplotypes. Genes listed below contain non-synonymous variants (ie. amino acid changes, regulatory mutations or splicing mutations) consistent with the identified singly causal haplotype (NOD for *Tip5;* CAST for *Tip8*). *Hip42* displayed a more complex haplotype pattern (WSB/NOD vs AJ/B6/NZO), and candidate selection is discussed in the main text. Genes with missense or nonsense variants (denoted by *).

| ***Tip5*** | ***Tip8*** | | ***Hip42*** |
|---|---|---|---|
| Ctsz | Fxyd5* | Siglecg | Ank3 |
| Tubb1 | Fxyd1 | Nkg7 | Cdk1 |
| Atp5e | Lgi4 | Cd33* | Tmem26 |
| Prelid3b | Fxyd3 | Siglece* | Slc16a9 |
| Zfp831* | Hpn | Klk13 | Fam13c |
| Edn3 | Scn1b | Klk8 | Rhobtb1 |
| Gm14391* | Gramd1a* | Klk7* | |
| Gm6710 | Pdcd2l* | Klk1b9* | |
| Zfp931 | Gpi1 | Klk1 | |
| | 4931406P16Rik | Clec11a | |
| | Kctd15 | Shank1 | |
| | Chst8 | Syt3 | |
| | Pepd | Lrrc4b | |
| | Cebpa | Josd2 | |
| | Slc7a10 | Spib | |
| | Lrp3* | Pold1 | |
| | Rhpn2 | Napsa* | |
| | Faap24 | Kcnc3 | |
| | Tdrd12* | Myh14 | |
| | Ankrd27* | Atf5 | |
| | Pdcd5 | Il4i1 | |
| | Dpy19l3 | Pnkp* | |
| | Tshz3* | Ptov1 | |
| | Ccne1 | Fuz | |
| | 1600014C10Rik | Tsks | |
| | Plekhf1 | Cpt1c* | |
| | Vstm2b | | |
| | Zfp975* | | |
| | Zfp715* | | |
