## Supplementary figures and images for "Host-pathogen genetic interactions underlie tuberculosis susceptibility"

### FigureS1_traitcorrplot.png

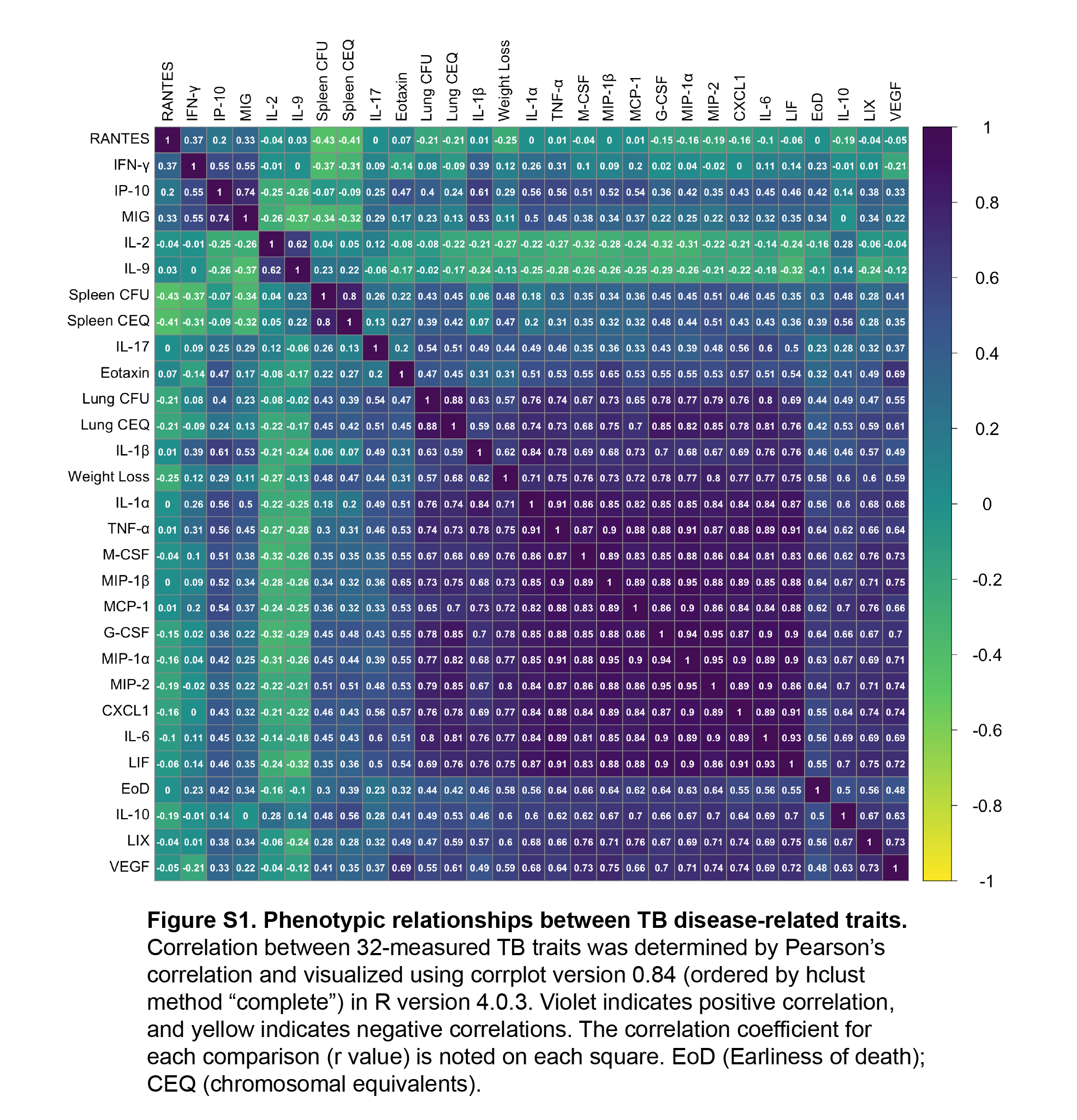

### FigureS2_QTLvisual.png

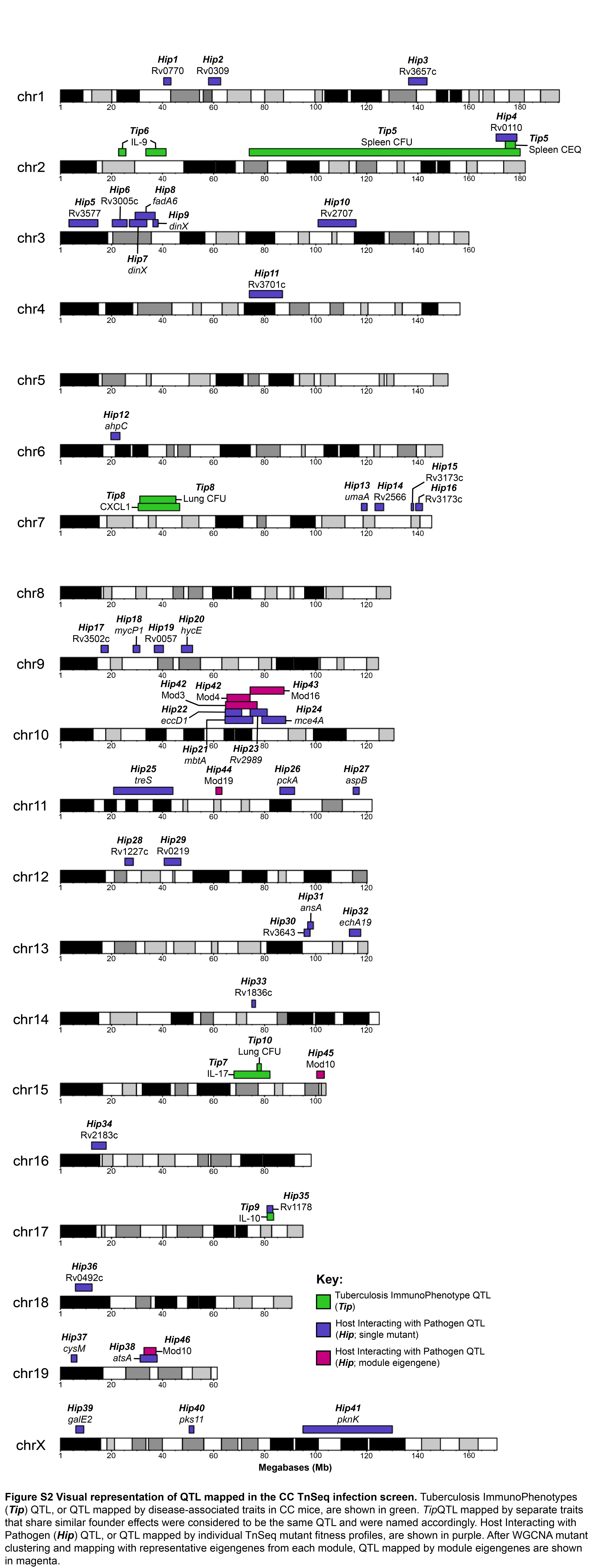

### FigureS3_ModuleTraitAssociation.png

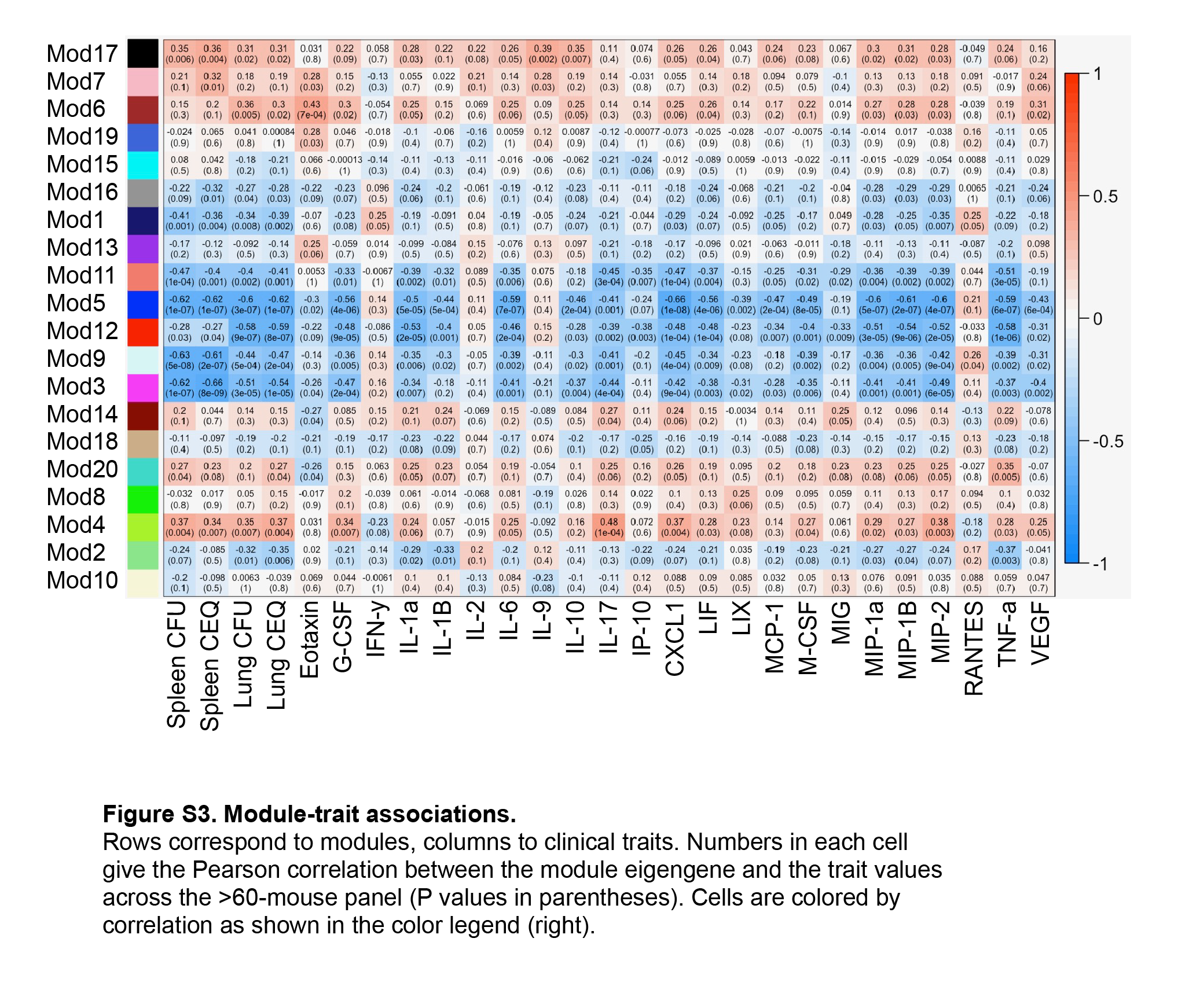
